# How much the response of rice genotypes to water deficit at the reproductive phase is dependent from environmental conditions?

**DOI:** 10.1101/2021.09.01.458547

**Authors:** Isabela Pereira de Lima, Tanguy Lafarge, Adriano Pereira de Castro, Sandrine Roques, Armelle Soutiras, Anne Clément-Vidal, Flávia Barbosa Silva Botelho, Marcel de Raïssac

## Abstract

Rice crop is known as particularly sensitive to water deficit, especially during the reproductive phase when growth of vegetative organs and formation of spikelets are simultaneous. Many works have focused on the response of rice plants to water deficits varying in timing, duration and intensity. Oppositely, the impact of the environmental conditions on the response to a given water deficit remains largely unknown. In order to test it, two experiments under contrasted conditions of temperature, radiation and VPD were conducted on six genotypes in greenhouse in Brazil (S) and in growth chamber in France (GC). The plants were submitted to the same mild water deficit at the reproductive phase, by adjusting FTSW at 0.4. Under irrigation, plant growth rate was reduced and crop duration extended in GC in relation to S: ultimately, this trade-off resulted in similar plant height and biomass in both environments. Under water deficit and in both environments, elongation rate decreased and was associated with an increase in soluble sugars in stem and flag leaf, while starch was reduced in S and negligible in GC because of the low radiation. This common biochemical response displayed a large gradient of values across environments and genotypes, but differentially impacted the branch and spikelet formation on the developing panicle: in carbon limiting conditions (GC), the increase in soluble sugars was associated with the reduction in branch and spikelet number, and conversely in S. At the morphological level, the maintenance of spikelet number on the panicle was correlated with the maintenance of flag leaf width in all genotypes and conditions, that was discussed according to the maintenance of the apical meristem size. Genotypes were discriminated and the study underlined the global tolerance of Cirad 409 and sensitivity of IAC 25.

## Introduction

Rice (*Oryza sativa* L.) is known to be particularly sensitive to soil water deficit because of its semi-aquatic origin (Kumar et al. 2014). Drought is therefore considered as the most important constraint reducing yield in rainfed areas (Kumar et al. 2008; Serraj et al. 2009). It impacts rice yield components according to its timing, duration and intensity: i) during the vegetative phase, development rate, plant height, leaf area and tillering are the most sensitive traits; ii) during the reproductive phase, panicle branching, spikelet formation and pollen viability may be critically damaged and; iii) after flowering, drought may jeopardize grain setting and filling with direct consequence on grain number and grain weight.

In Brazil, large areas are cultivated in upland rice in the central Cerrado and are submitted to drought. Heinemann et al. (2015) classified Brazilian upland ecosystems in three target populations of environment (TPE): a highly favorable environment (HFE) mainly stress-free; a favorable environment (FE) where mild water deficit occurs equally at reproductive and terminal phases and; a least favorable environment (LFE), dominated by more severe water deficit occurring at the reproductive phase. They have concluded on the priority for Brazil to improve upland rice tolerance to water deficit at the reproductive phase.

Literature converges in pinpointing the reproductive phase as the most sensitive one. According to Matsushima (1966), the period of time between minus eleven and minus three days before heading is the most sensitive period by its impact on final yield. In the same line, comparing the effect of water deficit at different development stages, Lilley and Fukai (1994) and Boonjung and Fukai (1996a) demonstrated that drought has non-significant or poor impact on rice subsequent development and grain yield when it occurs at vegetative stage, while the same deficit at reproductive phase causes yield and spikelet number reduction by 20-70 % and 60 %, respectively. This can be accounted for by the phenotypic plasticity of cereals at early stages because of the trade-offs between field panicle density, grain number per panicle and single grain weight (Wey et al. 1998; Siband et al. 1999). These trade-offs are highly expressed in rice in which tillering is more effective than in any other cereals.

With panicle initiation (PI) on the main tiller, plant enters the reproductive phase and faces a high demand in carbon, with the concomitance of the on-going leaf growth, the start in internode elongation and, to a lesser extent, the reproductive structure formation (Counce et al. 2000, Figure 1). At plant level, young tillers may emerge and act as a sink for carbon, before becoming autotrophic (Matsuo and Hoshikawa, 1993). At tiller level, demand in assimilate is high for the last four growing leaves and especially for the stem and peduncle elongation. Complete expansion of these organs is critical as: i) the three last leaves are responsible for up to 86 % of assimilate importation into the rice panicle under non limiting conditions (Cock and Yoshida, 1972), and up to 95 % into the wheat ear (El Wazziki et al., 2015) and; ii) the stem and peduncle elongation drives the achievement of panicle exertion and flowering ability. Any reduction in source leaf dimensioning may reduce assimilate supply to stem elongation and panicle formation. Oppositely, if the source is efficient, assimilate surplus can also be stored in stems and sheaths, in order to support efficiently grain filling in case of post-flowering water deficit (de Raïssac, 1992). The period between PI and flowering is thus a time of high competition for carbohydrates within a tiller, between the growth of existing vegetative organs and the initiation/differentiation/expansion of new reproductive structures, driving (i) the dimensioning of the future carbon source (leaf area); (ii) the panicle exertion ability (internode and peduncle length); and (iii) the dimensioning of the ultimate carbon sink (number of branches and spikelets).

**Figure 1.**
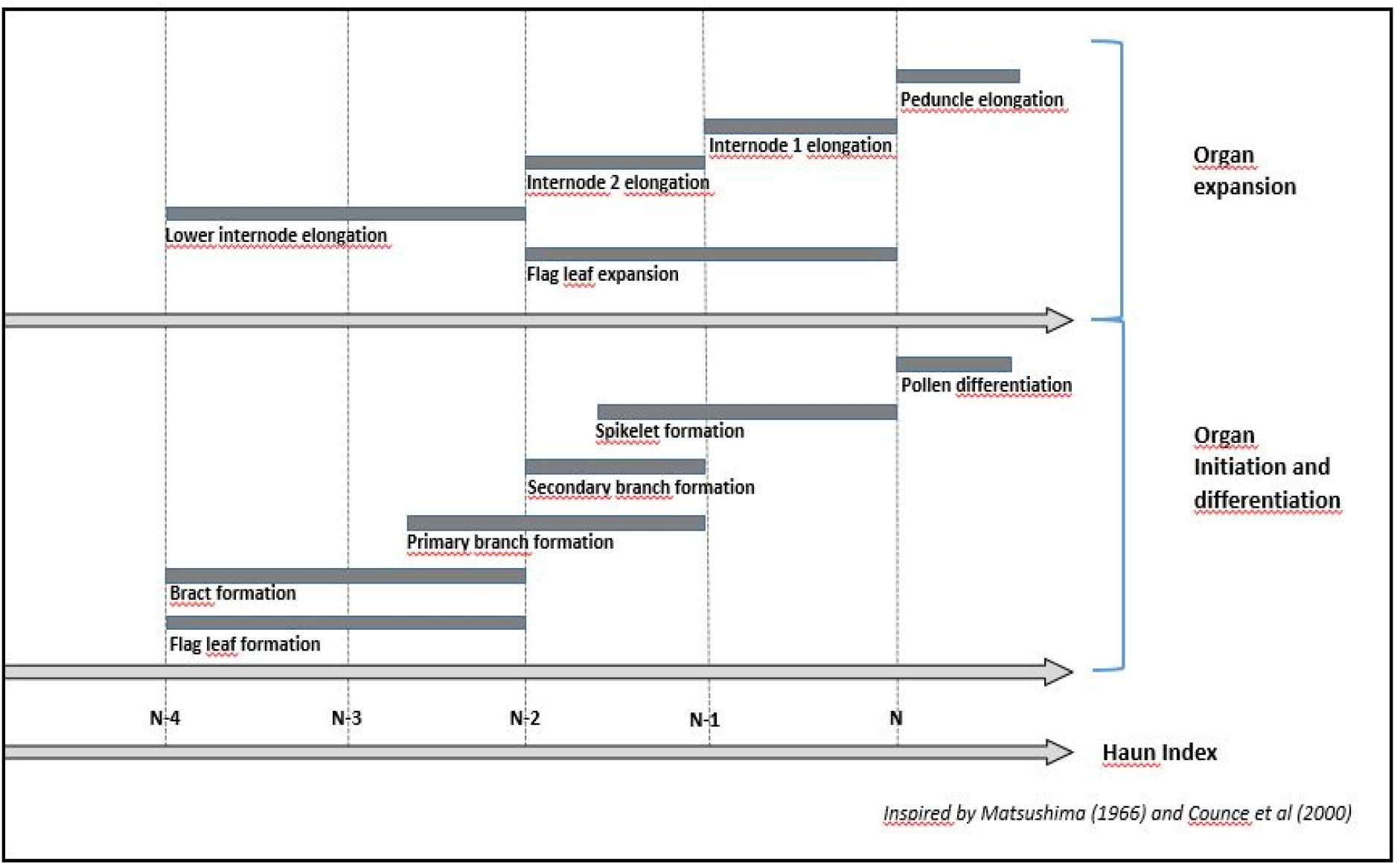
Timing of developmental processes during the reproductive phase in rice. The timing and duration of the sequential processes occuring after panicle initiation are represented according to Matsushima (1966) and Counce et al (2000), in relation with the Haun Index. N ref the time at flag leaf ligualtion (Haun Index = N) and N-4 the time at panicle initiation, given that it occurs four phyllochrones before flag leaf ligulation. Organ differentiation refers to the period whe is initiated and when first stages of its development take place (cell multiplication, tissue differentiation). Organ expansion refers to the generally rapid phase of expansion, once differentiation had set and as cell division and elongation are active.

With the occurrence of water deficit during the reproductive phase, tillering decreases or -if maintained-generates mainly unfertile tillers (Alou et al. 2018); the plant growth rate is slowed down, reducing grain number and potential grain size (Lilley and Fukai 1994a). In addition, flowering is systematically delayed (Boonjung and Fukai 1996b; Lilley and Fukai 1994a), up to 2 to 3 weeks depending on the intensity of the stress (Fischer and Fukai, 2003; Lafitte et al. 2003). Garrity and O’Toole (1994), and Swamy et al. (2017), found a negative correlation between the delay in flowering and yield, and proposed to use the delay as an indicator of plant sensitivity to drought (Subashri et al. 2009; Swamy et al. 2017). In some cases, the inflorescence does not emerge or emerges partially (Lafitte et al. 2003), preventing or reducing flowering. This delay and the poor panicle exertion can be associated with the lengthening in phyllochron and be the direct consequence of the decrease in peduncle elongation (Serraj et al. 2009; He and Serraj 2012), that accounts for 70-75% of spikelet sterility (O’Toole and Namuco, 1983). The reduction in branch and spikelet number, which is the main component driving the yield decline, is another major effect of the reproductive water deficit (Lilley and Fukai 1994b; Boonjung and Fukai 1996a). When the deficit occurs just before heading, it also affects pollen viability (Fig.1) and subsequently increases the spikelet sterility (Fischer and Fukai, 2003).

Under drought the question also raises whether growth is limited by leaf CO^2^ assimilation or by the intrinsic capacity of expansion in the growing tissues, that is to say whether growth is source or sink limited. Anciently comparing maize, soybean and sunflower, Boyer (1970) demonstrated that leaf enlargement decreases prior to photosynthesis in response to the reduction in water potential. This leads to an accumulation of sugars, as observed in sunflower (Dosio et al. 2011), in coffee tree (Franck et al. 2006) or in rice (Luquet et al. 2006; Luquet et al. 2008; Rebolledo et al. 2012), giving evidence that growth is sink - rather than source- limited, more precisely by the water status in the growing tissues rather than by the carbon availability (Muller et al. 2011; Tardieu et al. 2018). Pantin et al. (2011) and Pantin et al. (2012) even proposed a switch from a metabolic to a hydro-mechanical limitation of leaf growth during the course of leaf ontogeny. But beyond the leaf, knowledges are lacking for the inner sink organs, as internode, peduncle and panicle, that are heterotrophic tissues whose growth is protected from dehydration and dependent on plant C availability. With the pending question on the relationship between vegetative organ expansion and reproductive organ formation: is it synergistic or antagonistic?

The high complexity of combining adaptive with productive traits in the same genetic materials slowed down the genetic improvement for drought tolerance (Kumar et al. 2008). Breeding programs for tolerance opened two complementary ways. The first one relies on the direct selection through grain yield under drought, assuming that yield is the target objective beyond the sequential mechanisms that generated it. Works largely focus on the detection of QTL for yield and yield components (Lanceras et al. 2004; Kumar et al. 2008; Serraj et al. 2011; Kumar et al. 2014, Kumar et al. 2018; Sandhu et al. 2019;). The second one relies on “secondary traits”, that are defined as stable traits correlated to high yielding genetic materials in favorable conditions and in predominant stress situations (Lafitte et al. 2003). These “independent component traits associated with crop productivity” (Subashri et al. 2009) are indicators of tolerance (Sellamuthu et al. 2011; He and Serraj 2012). Ultimately, the “drought tolerance” *per se* concept remains unclear and could be questioned as drought specifically impacts plant development and yield according to its timing, its duration and its intensity: a mild water deficit at tillering shall not involve the same tolerance mechanisms than a severe one at reproductive phase.

In the present paper, we tested a set of six genotypes grown under two contrasted environments and submitted to a water deficit at reproductive phase. The objective is to explore whether, beyond the specific patterns of growth and development observed in the two situations, some generic traits can be highlighted as drought tolerance responses independent from environmental conditions.

## Material and Methods

### Genetic materials

Six genotypes were used within two experiments. They were extracted from the PRAY japonica collection, retrievable on line (GRiSP- Global Rice Phenotyping Network, 2020). The selection was made in order to compare short cycle materials with a large genetic diversity and distinct geographic origins (Table 1). Data collected in previous experiments in Brazil and in controlled environments in France (Table 2) show that the genotype classification for cycle duration is only partially maintained between experiments, with an approximatively two week lengthening of duration in France.

**Table 1.**
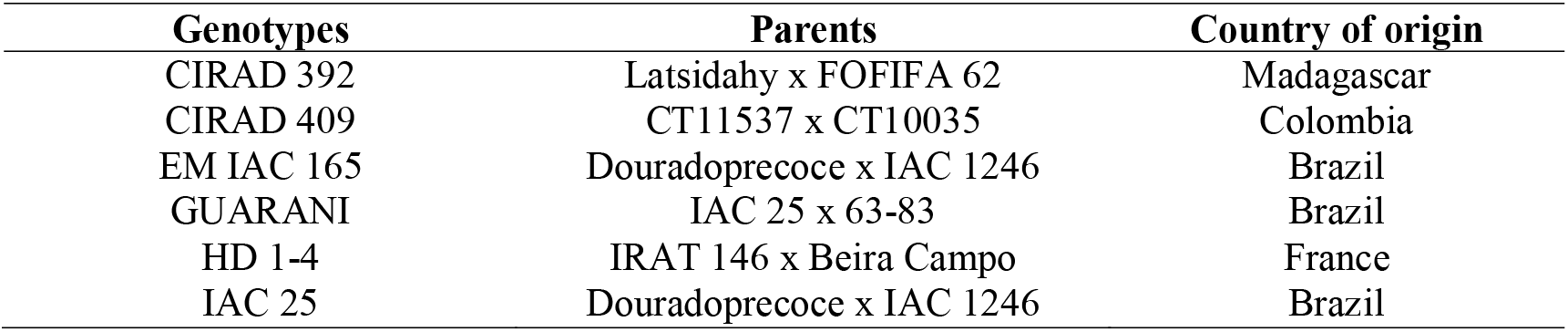
Upland rice genotypes used in the trials Sitis and Growth Chamber.

**Table 2.** Duration in days from planting to flowering obtained from previous experiments.

| **Genotypes** | **Days to Flowering** |  |  |
| --- | --- | --- | --- |
|  | **Field 2014,**<br>**Brazil** | **Growth Chamber**<br>**2016, France** | **Greenhouse**<br>**2008, France** |
| CIRAD 392 | 58 | 75 | 71 |
| CIRAD 409 | 56 | 69 | 70 |
| EM IAC 165 | 61 | 79 | 71 |
| GUARANI | 58 | 71 | 77 |
| HD 1-4 | 59 | 68 | 77 |
| IAC 25 | 59 | 73 | 71 |

### Sitis experiment (S)

This experiment was conducted in Goiânia, Brazil, at Embrapa Rice and Beans, at an altitude of 823 m, latitude 16°28’00”S and longitude 49°17’00”W. The SITIS (Integrated System for Induced Treatment of Drought) phenotyping platform was used. The platform, set up in the greenhouse, is composed of a set of 100 cm high and 25 cm diameter PVC pipes, each one filled with soil and placed upon an automatic scale that allows a continuous weighting of the pipes and monitoring of irrigation. The soil was a red latosol with medium texture, previously homogenized with a 1.25 cm mesh sieve to remove larger aggregates. The six genotypes were placed in a complete randomized block design with three replications and two water treatments: fully irrigated (IRR) and water stressed (STR) treatments, totalizing 36 plots. To secure three plants in each pot, a germination test has been carried out and plants were sown in excess on September 24, 2015. An application of 4 g of fertilizer NPK 04-14-08 was performed at planting, according to the recommendations for rice cultivation in relation with soil mineral analysis. Ten days after plant emergence, thinning was done to obtain three plants per pot. Climate conditions were monitored by an AKSO® device placed in the center of the greenhouse, continuously measuring air temperature and humidity, as well as solar radiation (Table 3).

**Table 3.**
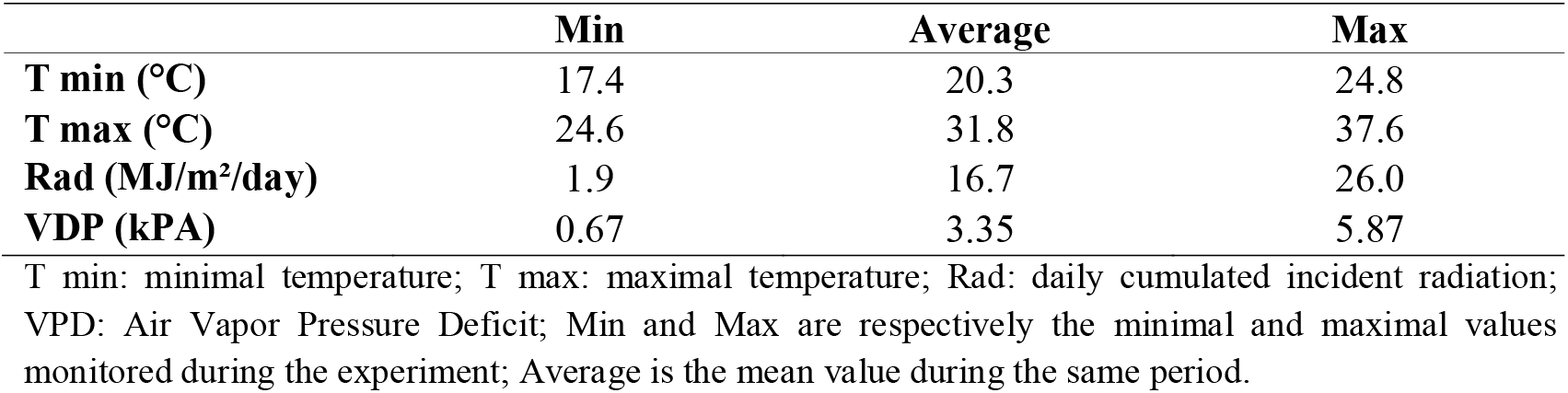
Climate conditions in Sitis experiment.

Soil physical analysis provided values of 24.5% and 11.5% for field capacity (FC) and wilting point (WP) respectively. Before planting, pots were slightly over-irrigated. The water in excess drained off and the weight at field capacity was determined 24 hours after this irrigation. Taking into account PVC pipe and plate weights, soil weights at FC and WP were then calculated. To quantify water availability within the pots, the Fraction of Transpirable Soil Water (FTSW) was calculated as the ratio between the actual available water and the maximum available water (Sinclair and Ludlow 1986), according to the following formula:

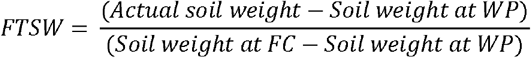

The pots of both treatments were adjusted daily to field capacity during the whole vegetative phase and the beginning of the reproductive phase. Two days before water stress application in the STR treatment, all pots were adjusted to FTSW = 0.8. The water stress was applied 15 days after the date of panicle initiation (PI), estimated from the 2014 Brazilian field experiment data (Table 2). Stress application started on October 30, 2015, for Cirad 392, Cirad 409 and Guarani, and on November 03, 2015 for EM IAC 165, HD 1-4 and IAC 25. After a 4-5 day period of dry-down, pots were daily adjusted to FTSW 0.4 with a bottom-up irrigation: water was simply provided in the plate below the pot, after observations had confirmed the presence of few primary roots at the base of the pot for each genotype, meaning that the whole soil profile was colonized. Water deficit treatment ended when at least 50% of main-tiller panicles of the irrigated treatment has emerged, that is to say when at least 5 plants on the 9 plants in the 3 irrigated replications were at heading stage.

In each pot, one plant tagged with a wool yarn and noted “plant 1” was observed for non-destructive phenological traits up to the end of the differential water treatments. This plant was then dissected while the two other plants of the pot were further grown at full irrigation from heading to grain maturity and harvesting (results not presented here).

### Growth Chamber experiment (GC)

Two experiments were carried out under the same climate conditions in a growth chamber in Montpellier in 2016 and 2017: it was set up as a 12/12 hours photoperiod, with day and night temperature and air humidity regulated at 28°C and 20°C, and at 65% and 90%, respectively (Table 4).

**Table 4.**
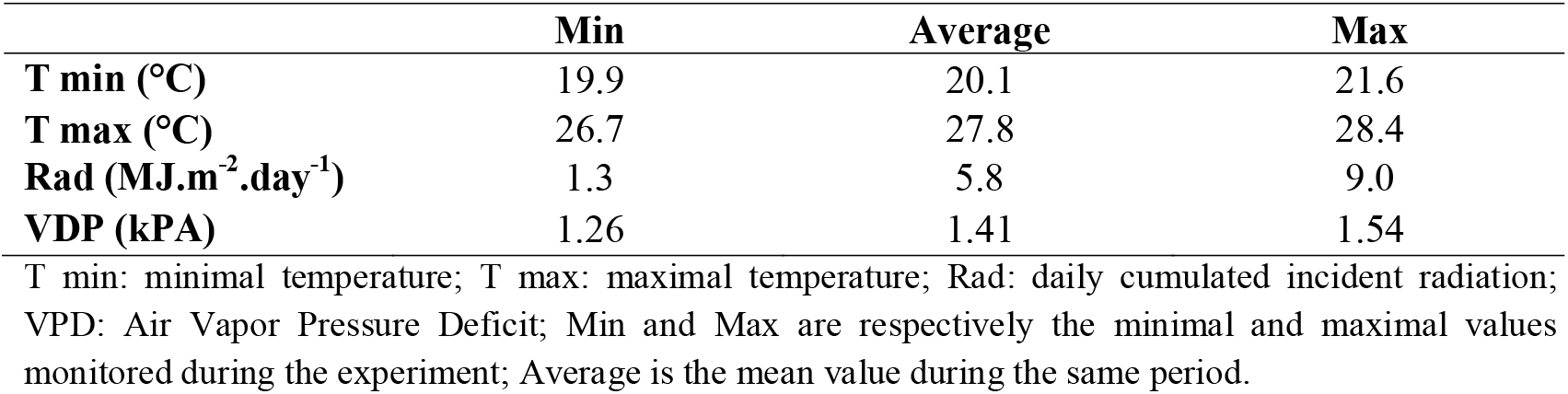
Climate conditions in Growth Chamber experiment.

|  | **Min** | **Average** | **Max** |
| --- | --- | --- | --- |
| **T min (°C)** | 19.9 | 20.1 | 21.6 |
| **T max (°C)** | 26.7 | 27.8 | 28.4 |
| **Rad (MJ.m<sup>-2</sup>.day<sup>-1</sup>)** | 1.3 | 5.8 | 9.0 |
| **VDP (kPA)** | 1.26 | 1.41 | 1.54 |

T min: minimal temperature; T max: maximal temperature; Rad: daily cumulated incident radiation; VPD: Air Vapor Pressure Deficit; Min and Max are respectively the minimal and maximal values monitored during the experiment; Average is the mean value during the same period.

A first methodological pre-experiment addressed the choice of soil substrate and the intensity of water deficit, as well as checking genotype cycle duration from planting to flowering. It was a complete randomized design with six genotypes and one water treatment (full irrigation). After germination, planting was achieved on 20 September, 2016. The number of days to panicle initiation was estimated by using the Haun index method (see below). Duration from planting to estimated panicle initiation varied between genotypes, from 26.4 days in HD 1-4 to 36.3 days in EM IAC 165. The reference genotype IR 64 was added, to which two levels of water deficit were tested from 5days after panicle initiation to panicle emergence: one adjusted each two days to FTSW 0.4, the other to FTSW 0.2, meaning that FTSW was lower than its ceiling value between two consecutive irrigations. The FTSW 0.2 treatment was finally too severe as it prevented any development of the plants during the reproductive phase. The FTSW 0.4 treatment was then selected, as a “mild” water deficit, equivalent to the one used in Sitis experiment.

The second and main experiment was conducted in a completely randomized design with six genotypes grown under two water treatments, irrigated (IRR) and stressed (STR), in three replications for a total of 36 pots. Each replicate was composed of a 3.5 liter polyethylene pot containing one single plant. Seeds were pre-germinated in an incubator set up at 30°C for three days. Three seedlings were transplanted on 5 January, 2017 and subsequently thinned to one plant per pot. The pots were filled with 1470 grams of a commercial referent substrate (Tref Riz Cirad 2), specifically adapted to rice, mixed with seven grams of Basacote 6M+, a slow-release fertilizer. For each pot, a soil sample was weighted and oven-dried in order to determine the initial effective soil humidity and the real soil dry weight. Pot weights at FC and at WP were then deduced in order to calculate FTSW. The water deficit treatment was applied by withholding water supply 5 days after the date of panicle initiation estimated from the pre-experiment. At the start of the water deficit application, the pots were covered with polystyrene micro balls to avoid water loss by direct soil evaporation, allowing the measurement of the plant water consumption between two consecutive irrigations. Water was supplied every two days by top irrigation in order to adjust to FTSW 0.8 and 0.4 in IRR and STR treatments, respectively. As in Sitis, the water deficit treatment was maintained in STR plants up to panicle emergence on the main tiller of the IRR plants. At this date, all plants were dissected and their samples dried in oven.

### Measurements in both experiments

#### Thermal time

Mean air temperature was calculated on a daily step in both experiments. Thermal time was then calculated using Samara model values (Kumar et al. 2017), with the following cardinal temperatures: Tb = 10, Topt1 = 28, Topt2 = 36, Tmax = 44. The cumulated thermal time from planting to heading increased quicker in S than in GC (Figure 2). The mean daily cumulated thermal time was 17.5 °C days and 13.7 °C days in S and GC respectively.

**Figure 2.**
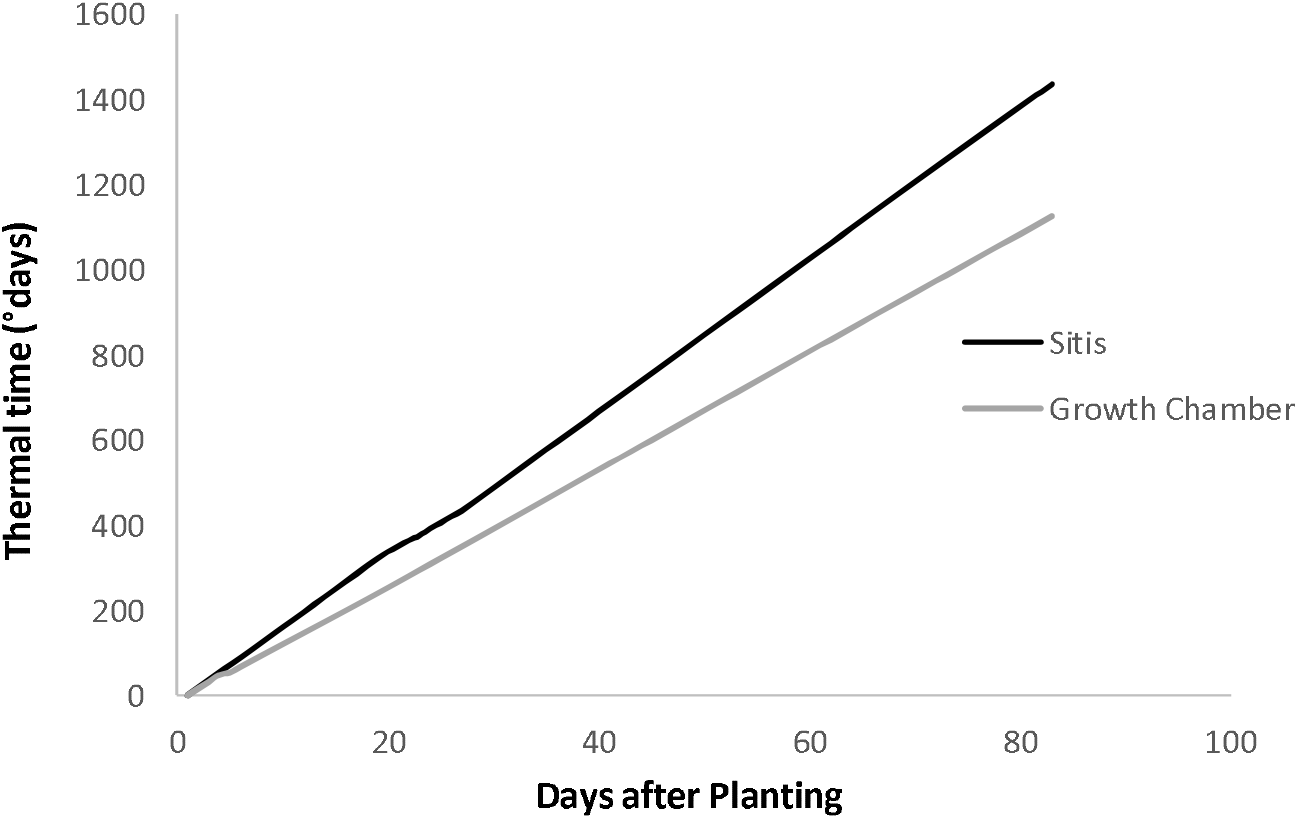
Cumulated thermal time in Sitis and growth chamber experiments in function of calendar time. Thermal time was calculated on an hourly step in GC and daily step in Sitis. Mean temperatures were calulated as (Tmax + T min)/2. We then used cardinal temperatures from Samara model : T_b_ = 10, T_opt1_ = 28, T_opt2_ = 36, T_max_ = 44 to calculate the cumulated thermal time at hourly and daily steps in GC and S respectively

#### Phenological development monitoring

The Haun Index (HI, Haun, 1973) was monitored 3 times a week on the main tiller of each plant. HI is calculated as the number of fully expanded leaves on the main stem plus the estimated ratio of current length to final length of the growing leaf. The phyllochron was calculated as the time -in days or thermal time- separating two successive fully expanded leaves on the main tiller. The early phyllochron was determined before PI and under non-limiting water supply for all genotypes, that is between 14 and 27 days after planting (DAP) in Sitis, and between 11 and 24 DAP in GC. Phyllochron was also determined from the onset of differentiated water treatments to the maximum Haun index (Max HI), when flag leaf expansion ended.

For each plant in a pot, the date of PI was retrospectively estimated, according to the method illustrated in Figure 3 for Cirad 409 grown in GC under irrigated conditions: because the final leaf number (LeafNb) on the main tiller is observed at panicle emergence and because PI is known to occur at LeafNb-4 (Nemoto et al. 1995), the precise date of PI was estimated by interpolation between two consecutive phenological observations circumscribing the target PI. In the illustrated case (Figure3), LeafNb is 10, PI consequently occurs at HI=6 and PI date estimated by interpolation at 34 DAP. This method was retrospectively applied to all individual plants in both experiments. Plant height, defined as the distance between the soil and the last ligule on the main stem, was measured twice a week. The heading stage was considered in each genotype when more than 50% of the main tiller panicles had emerged on the irrigated plants.

**Figure 3.**
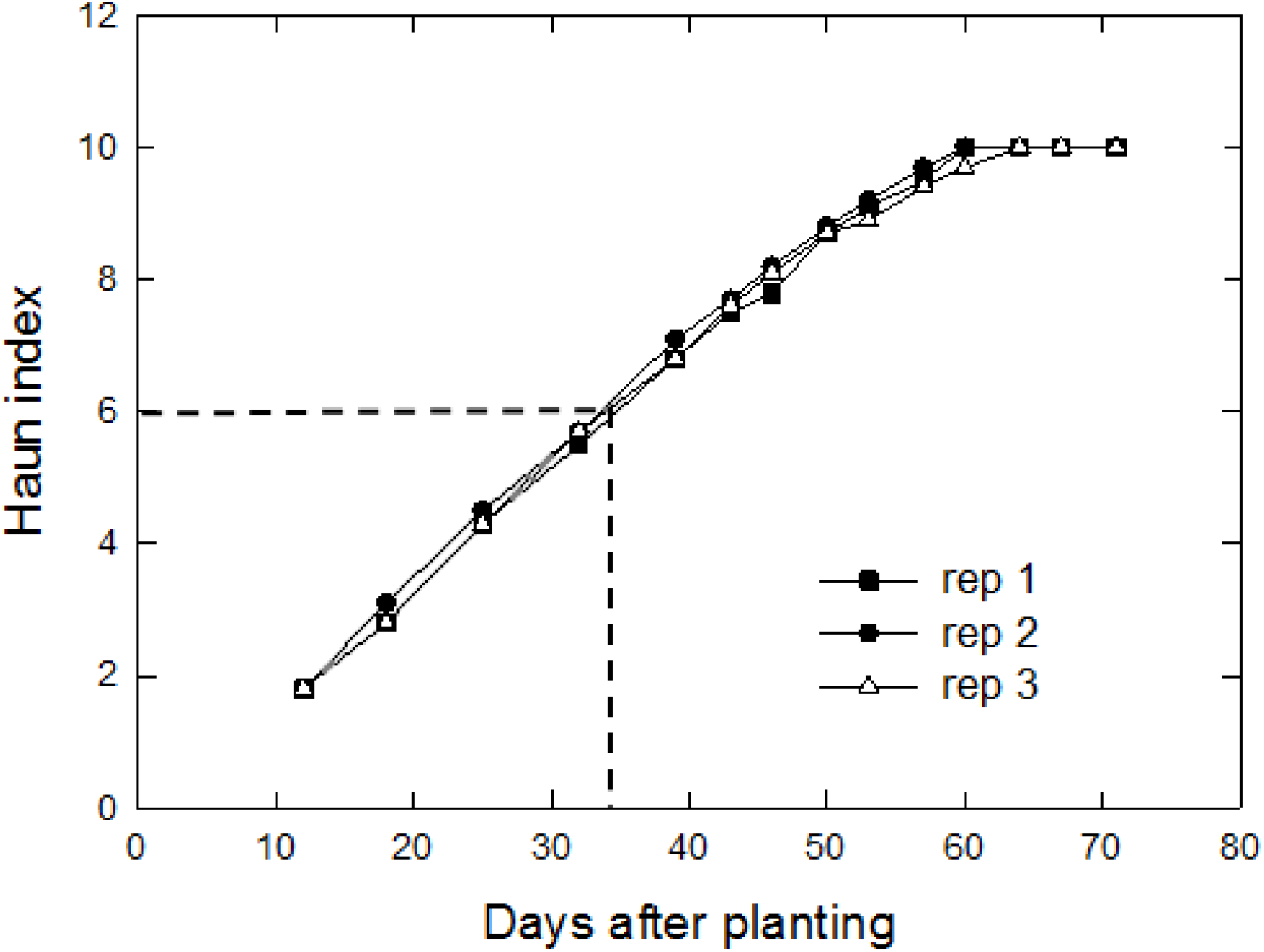
Determination of the panicle initiation date on irrigated Cirad 409 in the growth chamber experiment. For each replicate, the date of panicle initiation is determined as the ligulation time of the N-4 leaf, on S and GC experiments. In the present case study, the plant has developped 9 leaves on the main tiller; the panicle initiation occured at the ligulation of the 5th leaf, that is at HI = 5. Interpolation between the closest observations determines the PI date : 17/10/15. Days between panicle initiation and water deficit application was then calculated for each plant on both experiments.

At panicle emergence in both experiments, after tiller counting, plant dissection has been done, to determine the following data:

- Lengths and widths (diameters) of internode 1 and 2 on the main tiller, considering internode 1 as the first internode below the peduncle;
- Dry weights of panicle, leaves, internodes and nodes on the main tiller;
- Number of first and second order branches on the main tiller panicle;
- Number of spikelets on the main tiller;
- Dry weights of remaining tillers;

In addition, the length and width of all leaves on the main tiller were measured as soon as they were fully expanded. Only in S, plant dry weight was measured at thinning at 14 DAP. Only in GC, the panicle main axis length and the total length of primary and secondary branches were measured.

#### Carbohydrate analysis

Flag leaf blade (FL) and internodes 1 and 2 (IN1 and IN2) were sampled early in the morning, and plunged into liquid nitrogen right after sampling, freeze-dried (72h), grinded with a ball mill Retsch MM400 (particules < 50 μm) and then stored in a freezer at −80°C. Sugar content was measured in Sitis according to the method described by Gibon et al. (2009) and in the growth chamber by Luquet et al. (2006). The sugar extractions and starch hydrolysis are similar between both methods, the methodological difference concerns only the sugar quantification. The first method uses spectrophotometry and the second one is based on high performance liquid chromatography. Samples were previously used to ensure that the results could be comparable. The results are expressed as glucose equivalents per unit dry matter (mg.g^−1^) for starch (Starch) and as soluble sugar (SolSug) per unit dry matter (mg.g^−1^) corresponding to the amount in hexoses and sucrose.

#### Gas exchange measurement

In S, transpiration was measured at the end of the experiment, between 9 and 11 a.m., on the flag leaf of each plant n°1, using the LC-Pro-IRGA® device, regulated at a 300 ml.min^−1^ air flow with a 1200 μmol.m^−2^.s^−1^light source.

In GC, photosynthesis and transpiration were measured the day before dissection by the Walz GFS 3000.

#### Derived variable calculation

The main tiller elongation rate was calculated as the increase in height during the stress period (in IRR and STR plants) divided by the number of days. An estimation of the relative plant growth rate (RGR) was calculated differently in S and GC. In S, genotype shoot dry weight was measured on six plants from the IRR treatment at 14 DAP-at time of thinning- and at panicle emergence. RGR was then calculated for IRR treatment as following:

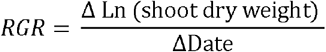

The mean genotype RGR was then used to estimate by intrapolation shoot dry weight of plants at the onset of water deficit application. Considering shoot dry weight at the onset of water deficit and at heading (end of the experiment), RGR (g.day^−1^) and GR (Growth rate - g.day^−1^) were calculated during the whole period when differential water treatments were effective. In GC, a strong allometric correlation between height and biomass was used to estimate the shoot dry weight of the plants at the onset of water deficit application.

In GC, because pots were covered by polystyrene bills preventing direct soil evaporation, the monitoring of pot weights before and after irrigation allows calculating the daily plant transpiration, as well as a cumulated plant transpiration during the differential water treatment period. The cumulated water use efficiency was then calculated as the ratio of the increase in biomass to the amount of water consumed during the same period.

To quantify genotype response to water deficit, response indexes were calculated as the relative variation between IRR and STR treatments, as defined below:

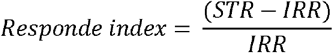

Where:

STR is the genotype adjusted mean of the considered variable for the water deficit treatment;
IRR is the adjusted mean of the considered variable for the irrigated treatment.

#### Statistical analysis

Anovas were run using R packages “agricolae” (de Mendiburu 2014), “ScottKnot” (Jelihovschi et al. 2014), “doBy” (Hojsgaard et al. 2019) and “ggplot2” (Wickham 2016). The response indexes were validated by a test of orthogonal contrasts, where the adjusted means of the IRR treatment were compared to the adjusted means of the STR treatment, using F Test and R software.

In order to compare traits measured on both experiments under irrigation, we conducted a split-plot analysis, considering they displayed a completely randomized design, with a first factor at two levels (sites) and a second factor at six levels (genotypes).

## Results

### Effect of environment on plant phenotypic traits under irrigated conditions

The rice plants grown under non-limiting water supply across both experiments displayed significant differences in their phenotypic traits, cumulating genetic and environment effects and, in most cases, interaction effects (Table 5). The vegetative phase (VegDuration) increased in average by 12,1 days from S to GC. In thermal time (Fig. 2), the increase was only 80°C.days, and could be accounted for by the increase in the average number of developed leaves on the main stem, from 8.9 to 10.4. Two genotypes, Cirad 392 and EM IAC 165, developped 2.6 and 2.3 extra-leaves respectively and delayed panicle initiation even more than the others. The reproductive phase (ReprDuration) was extended by 7 days in GC, without any interaction effect. But, in thermal time, the phase remained stable across sites, at 580 and 560 °C.days respectively. In other words, the thermal time required for the emergence of the four last leaves and the panicle was equal on both experiments.

**Tableau 5.**

|  | Significancy |  |  | Site Effect |  | Cirad 392 |  | Cirad 409 |  | EM IAC 165 |  | Guarani |  | HD 1-4 |  | IAC 25 |  |
| --- | --- | --- | --- | --- | --- | --- | --- | --- | --- | --- | --- | --- | --- | --- | --- | --- | --- |
| Trait | Site | Gtype | Site x Gtype | S | GC | S | GC | S | GC | S | GC | S | GC | S | GC | S | GC |
| VegDuration | *** | *** | *** | 22,4 a | 34,5 b | 22,0 a | 36,3 cd | 21,0 a | 33,3 bc | 22,3 a | 41,0 d | 22,6 a | 34 bc | 22,3 a | 30,7 b | 24,0 a | 31,7 bc |
| LeafNb | *** | *** | *** | 8,9 a | 10,4 b | 9,0 ab | 11,6 c | 8,7 a | 10,0 b | 9,0 ab | 11,3 c | 9,0 ab | 10,0 b | 8,7 a | 9,3 abb | 9,0 ab | 10,0 b |
| TillerNb | *** | * | ns | 9,7 b | 6,8 a | 11,0 c | 10,0 abc | 9,0 abc | 7,3 abc | 10,3 bc | 6,3 abc | 10,3 bc | 5,7 ab | 9,0 abc | 6,3 abc | 8,7 abc | 5,3 a |
| ReprDuration | *** | *** | ns | 33,1 a | 40,8 b | 34,0 ab | 43,7 e | 29,0 a | 36,7 bc | 34,7 b | 43,0 de | 33,3 ab | 38,0 bcd | 34,7 b | 41,3 cde | 33,0 ab | 42,3 de |
| TotBranchNb | ns | ** | *** | 35,2 a | 37,9 a | 23,0 a | 33,0 abc | 32,3 abc | 39,3 bcd | 42,7 cd | 51,7 d | 26,7 ab | 33,3 abc | 42,7 cd | 38,0 bcd | 44,0 cd | 32,0 abc |
| SpikeletNb | ns | ** | *** | 155 a | 156 a | 119 a | 133 ab | 141 ab | 145 ab | 180 bc | 217 c | 146 ab | 139 ab | 159 abc | 161 abc | 186 bc | 143 ab |
| SpkltFormRate | * | ** | *** | 10,5 b | 9,6 a | 7,4 a | 8,3 ab | 12,8 cd | 9,1 abc | 11,2 abcd | 13,6 d | 9,7 abcd | 8,7 ab | 10,6 abcd | 9,4 abc | 11,7 bcd | 8,4 ab |
| Phyllochron | ns | ns | ns | 7,8 a | 7,8 a | 8,2 a | 9,9 a | 8,7 a | 6,9 a | 9,2 a | 7,9 a | 7,3 a | 7,3 a | 7,5 a | 6,5 a | 6,2 a | 8,4 ab |
| FLLength | *** | ns | ns | 436 a | 604 b | 396 a | 583 ab | 441 ab | 615 ab | 511 ab | 607 ab | 421 ab | 591 ab | 394 a | 586 ab | 453 ab | 640 b |
| FLWidth | *** | *** | ns | 17,3 a | 18,6 b | 13,3 a | 15,0 a | 13,1 a | 15,3 ab | 19,1 c | 21,0 c | 18,3 bc | 19,6 c | 19,3 c | 20,3 c | 20,6 c | 20,7 c |
| IN1Length | ns | ns | ns | 166 a | 190 a | 180 a | 181 a | 151 a | 150 a | 188 a | 182 a | 162 a | 202 a | 142 a | 198 a | 174 a | 227 a |
| IN1Width | * | ** | ns | 5,0 a | 5,4 b | 5,0 ab | 5,6 b | 4,0 a | 4,6 ab | 5,6 b | 5,9 b | 5,0 ab | 5,0 ab | 5,3 ab | 5,6 b | 5,3 ab | 5,7 b |
| IN2Length | ns | *** | ns | 141 a | 140 a | 144 abc | 123 ab | 73 a | 80 a | 145 abc | 162 bc | 160 bc | 143 abc | 139 abc | 125 ab | 184 bc | 208 c |
| IN2Width | ** | *** | *** | 5,5 a | 5,9 b | 6,0 cdef | 6,4 ef | 4,0 a | 4,7 ab | 6,3 def | 6,8 f | 5,3 bcd | 5,1 bc | 6,0 cdef | 5,9 cdef | 5,8 cde | 6,5 ef |
| PedLength | ns | * | ns | 194 a | 199 a | 358 a | 237 a | 102 a | 126 a | 194 a | 155 a | 207 a | 264 a | 128 a | 146 a | 175 a | 268 a |
| PlantHeight | *** | *** | * | 697 a | 813 b | 699 bc | 717 bc | 564 a | 648 ab | 721 bc | 887 ef | 751 bcd | 854 def | 645 ab | 806 cde | 805 cde | 967 f |
| ElongRate | *** | *** | * | 26,3 b | 14,6 a | 25,8 de | 11,1 a | 26,1 de | 11,3 a | 25,0 cde | 14,0 a | 26,1 de | 16,4 ab | 24,4 cd | 14,8 ab | 30,4 e | 19,7 bc |
| GrowthRate | *** | *** | *** | 1,14 b | 0,40 a | 0,69 bc | 0,59 ab | 1,17 d | 0,26 a | 1,29 de | 0,29 a | 1,00 cd | 0,42 ab | 1,16 d | 0,35 ab | 1,52 e | 0,46 ab |
| TotBiomass | ns | *** | *** | 16,2 a | 15,7 a | 11,8 ab | 23,6 d | 12,8 ab | 8,5 a | 18,0 bcd | 15,5 abc | 17,0 bcd | 15,6 abc | 16,3 abcd | 13,7 abc | 21,3 cd | 17,3 bcd |
| LeafTransp | ns | ns | *** | 4,5 a | 4,4 a | 3,3 a | 5,0 ab | 3,3 a | 4,6 ab | 4,8 ab | 3,9 ab | 5,1 ab | 4,2 ab | 5,1 ab | 5,0 ab | 5,6 b | 3,8 ab |
| FLSolSug | ns | ns | ns | 65,9 a | 60,7 a | 70,3 a | 53,0 a | 72,0 a | 59,4 a | 60,0 a | 67,5 a | 72,6 a | 59,8 a | 52,7 a | 63,6 a | 67,9 a | 60,6 a |
| FLStarch | *** | * | * | 64,4 b | 0,6 a | 67,6 bc | 0,4 a | 73,1 c | 1,1 a | 60,0 b | 0,5 a | 62,2 bc | 0,5 a | 60,4 b | 1,0 a | 63,3 bc | 0,4 a |
| IN1SolSug | *** | *** | *** | 75,3 a | 130,9 b | 75,6 a | 186,5 c | 62,3 a | 61,1 a | 82,7 a | 164,8 bc | 74,6 a | 100,4 ab | 77,2 a | 110,4 ab | 79,1 a | 162,3 bc |
| IN1starch | *** | ** | ** | 264,0 b | 3,8 a | 321,8 d | 7,8 a | 307,7 d | 5,7 a | 276,3 cd | 2,7 a | 221,9 bc | 1,1 a | 195,9 b | 2,7 a | 260,5 bcd | 2,9 a |
| IN2SolSug | ns | ns | ns | 55,5 a | 78,6 a | 38,7 a | 131,0 a | 55,2 a | 53,8 a | 36,0 a | 77,1 a | 56,7 a | 60,8 a | 74,8 a | 53,9 a | 71,6 a | 95,0 a |
| IN2starch | *** | ns | ns | 126,9 b | 24,3 a | 118,6 cd | 36,8 ab | 104,0 bcd | 36,6 ab | 126,3 cd | 5,2 a | 142,2 d | 5,8 a | 129,4 cd | 59,5 abc | 140,9 d | 1,8 a |

The whole plant growth rate during reproductive phase (GrowthRate) decreased by 64 % from S to GC (1.14 and 0.40 g.day^−1^ respectively, Fig. 4a) which was associated with a 65 % decrease in the incoming radiation (from 16.7 MJ.m^−2^.day^−1^ to 5.8 MJ.m^−2^.day^−1^ respectively, Tables 3 and 4). Under S conditions, Cirad 392 (0.69 g.day^−1^) was characterized with the lowest value and IAC 25 (1.52 g.day^−1^) with the highest one. Low radiation in GC was also associated with a significant decline in tiller number per plant (from 9.7 to 6.8). Due to the delay in heading time in GC, the final whole plant biomass was globally not significantly different between both experiments (16.2 g in S and 15.7 g in GC), but GxE interactions were highly significant (Fig. 4b): in the case of Cirad 392, a genotype improved for cold tolerance on the Malagasy plateau, its biomass was the lowest in S while it was the highest in GC (Fig.4b).

**Figure 4.**
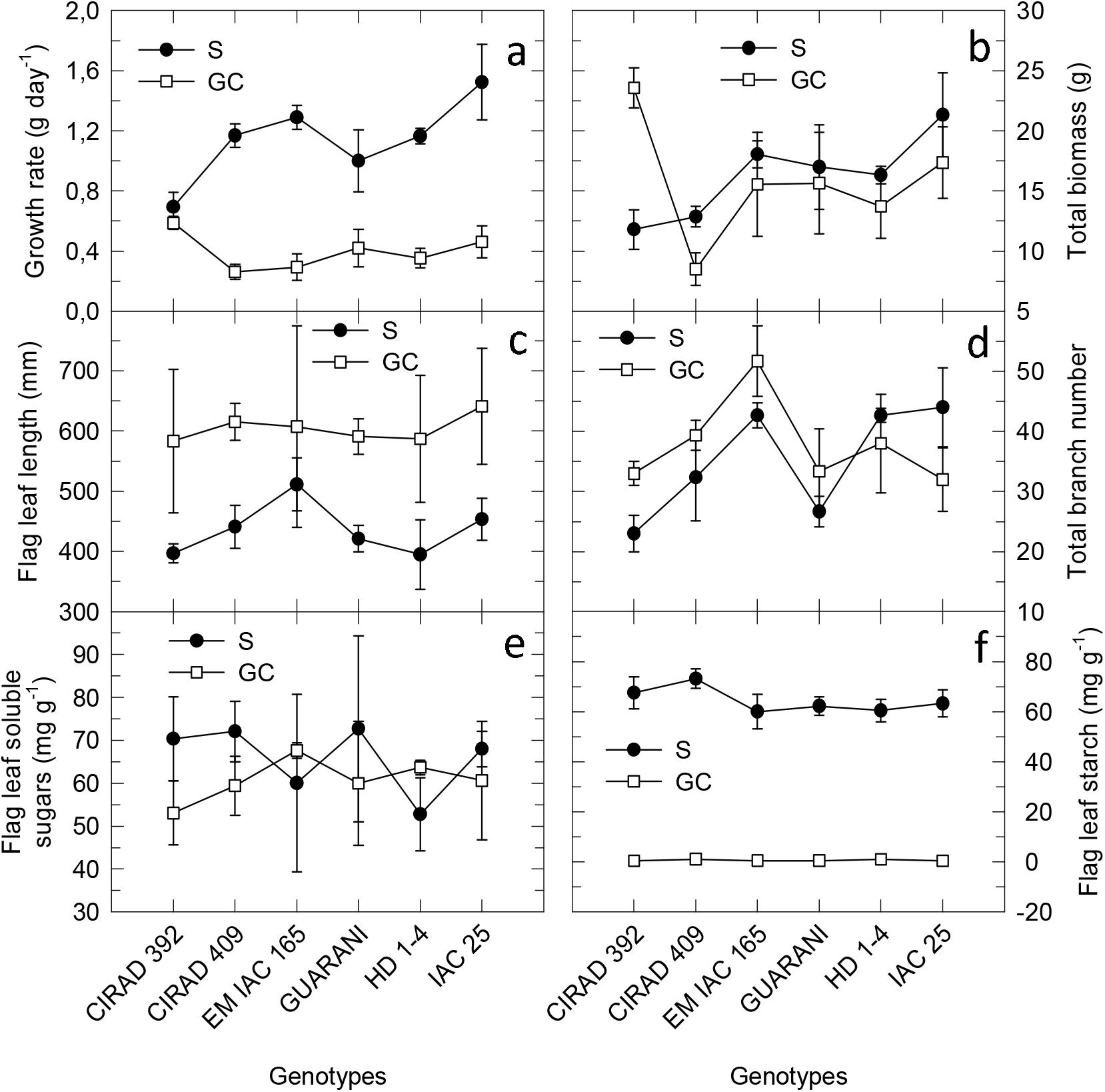
Phenotypic traits under irrigated conditions in both sites. S = Sitis greenhouse platform in Brasil; GC = Growth Chamber in France

Organ size was systematically larger in GC than in S (Table 5): the main stem was taller (813 vs 697 mm); flag leaf was longer (604 vs 436 mm, Fig. 4c) and wider (18.6 vs 17.3 mm); internodes were wider (5.4 vs 5.0 mm for IN1, 5.9 vs 5.5 mm for IN2). In S, the reported reduction in leaf size occurred in conditions where VPD was significantly higher: its higher elongation rate (26.3 mm. day^−1^ vs 14.6) did not compensate the reduction in growth duration under high VPD. Concentrations in soluble sugar varied in the same range across both environments in FL (66.0 mg.g^−1^ in S vs 60.7 mg.g^−1^ in GC, Fig. 4e) and in IN2 (55.2 mg.g^−1^ in S vs 78.4 mg.g^−1^ in GC). In contrast, concentrations in starch were drastically reduced from S to GC in these 2 organs (from 54.5 mg.g^−1^ to 0.7 mg.g^−1^ in FL, Fig. 4f, and from 128.4 mg.g^−1^ to 25.4 mg.g^−1^ in IN2) which was associated with the weakness of C source in GC characterized with low irradiance, that might have prevented the plant to store extra-assimilates (Fig. 4f).

The formation of the reproductive structure was poorly affected despite the significant shift in development patterns from one site to the other: the branch number (TotBranchNb) and the spikelet number (SpikeletNb) on the panicle of the main stem were not significantly modified: 35.2 vs 37.9 for TotBranchNb (Fig. 4d) and 155.2 vs 156.7 for SpikeletNb, in S and GC respectively. Nevertheless, interactions were highly significant and genotypes differed in their response to the environment: low performance was reported with IAC 25 in GC conditions and in Cirad 392 in S conditions (Fig. 4). Finally, the transpiration rate of the flag leaf (LeafTransp) measured right before heading did not significantly differ between experiments (Table 5).

### Impact of water deficit timing on developmental processes

The *a posteriori* determination of the actual occurrence of water deficit in relation with panicle initiation was done at the individual genotype basis. As described in Material and Methods, the date of PI was retrospectively estimated for each individual plant. The time lag between PI and the actual start of water deficit was determined. Then, a global index of the reproductive phase progress at time of water deficit application was calculated as the ratio in days of the elapsed time from PI to the onset of water deficit to the total reproductive phase duration (Table 6).

**Table 6.**
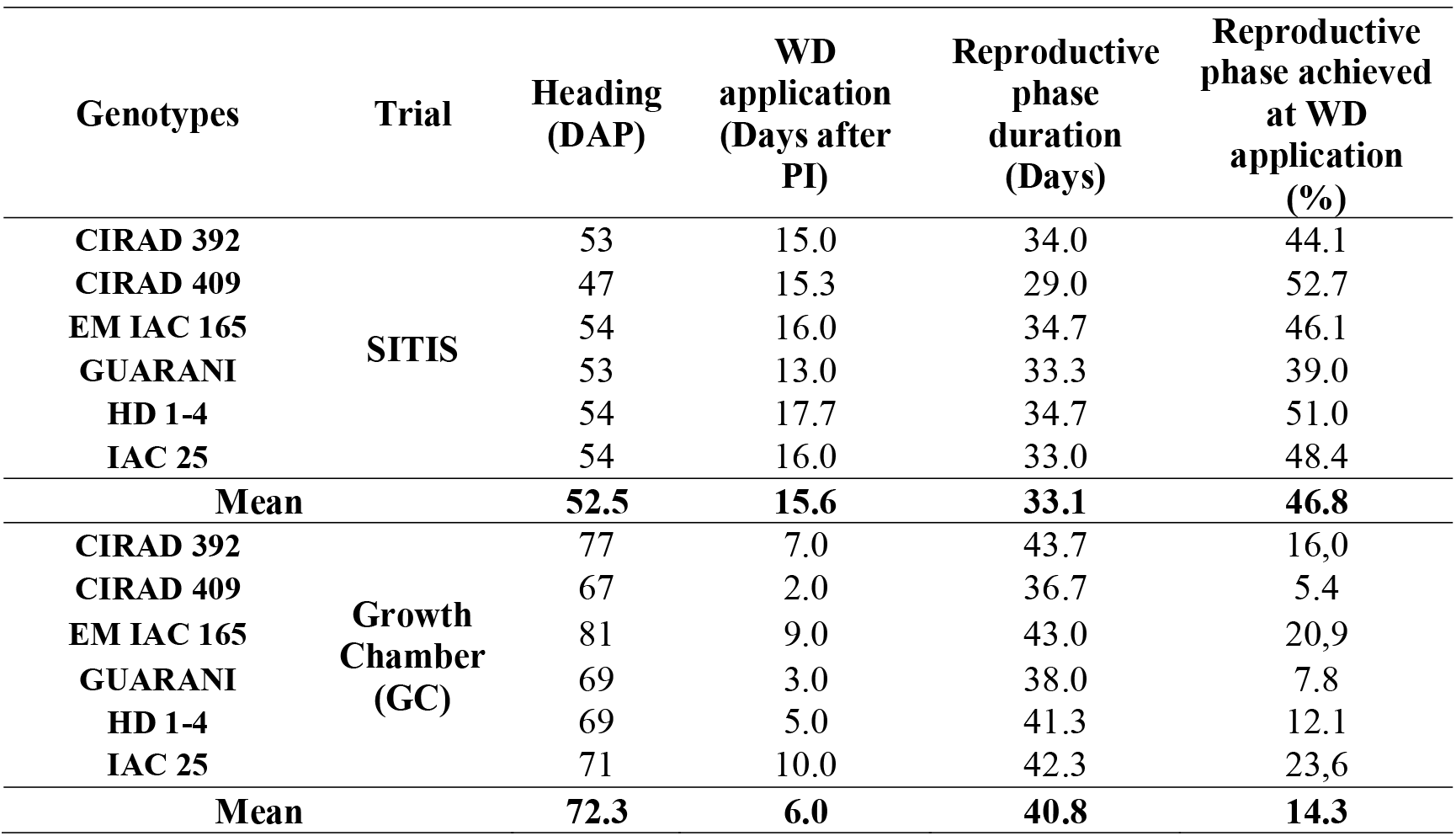
Mean progress of reproductive phase at start of water deficit application.

| **Genotypes** | **Trial** | **Heading**<br>**(DAP)** | **WD**<br>**application**<br>**(Days after**<br>**PI)** | **Reproductive**<br>**phase**<br>**duration**<br>**(Days)** | **Reproductive**<br>**phase achieved**<br>**at WD**<br>**application**<br>**(%)** |
| --- | --- | --- | --- | --- | --- |
| **CIRAD 392** | **SITIS** | 53 | 15.0 | 34.0 | 44.1 |
| **CIRAD 409** |  | 47 | 15.3 | 29.0 | 52.7 |
| **EM IAC 165** |  | 54 | 16.0 | 34.7 | 46.1 |
| **GUARANI** |  | 53 | 13.0 | 33.3 | 39.0 |
| **HD 1-4** |  | 54 | 17.7 | 34.7 | 51.0 |
| **IAC 25** |  | 54 | 16.0 | 33.0 | 48.4 |
| **Mean** |  | **52.5** | **15.6** | **33.1** | **46.8** |
| **CIRAD 392** | **Growth**<br>**Chamber**<br>**(GC)** | 77 | 7.0 | 43.7 | 16,0 |
| **CIRAD 409** |  | 67 | 2.0 | 36.7 | 5.4 |
| **EM IAC 165** |  | 81 | 9.0 | 43.0 | 20,9 |
| **GUARANI** |  | 69 | 3.0 | 38.0 | 7.8 |
| **HD 1-4** |  | 69 | 5.0 | 41.3 | 12.1 |
| **IAC 25** |  | 71 | 10.0 | 42.3 | 23,6 |
| **Mean** |  | **72.3** | **6.0** | **40.8** | **14.3** |

Water deficit (WD) occurred earlier in GC than in S with respect to the plant phenology, with actual dates very near from the target ones (Table 6): in average, its onset was 6.0 days after PI in GC and 15.6 days in S, with few variability across genotypes. At the genotype level, and according to the specific date of application and the reproductive phase duration, the relative progress of the reproductive phase at WD start was calculated as 14.3 % in GC, and 46.8 % in S.

The level to which each basic phenological process was affected due to the application of water deficit was finally computed for each genotype and in both experiments (Table 7 a and b), with reference to the timing and position of each one respective to the occurrence of WD (Figure 1). Growth processes occuring just after panicle initiation were not affected by water stress in S (Table 7a): this was the case for the first stages of flag leaf development and elongation of lower internodes of the main stem. Branching and elongation of internode 2 were partially affected, except with Cirad 409. Subsequent processes, including spikelet formation, pollen differentiation, elongation of internode 1 and peduncle were noted as fully affected by water stress (see below). In contrast in GC, the early processes, as flag leaf formation and elongation of lower internodes were partially affected (Table 7b), all other processes being noted as fully affected. The robustness of these assumptions were confirmed by comparing the flag leaf length across both experiments. In our estimation, the water deficit did not occur during the leaf formation in S while it did in GC. An anova conducted separately on both experiments confirmed this assumption, with a significant effect of WD in flag leaf length in GC only (Table 8 and Fig. 5). As no Treatments x Genotypes interaction was observed in GC, it is reported that leaf elongation across genotypes reacted the same way to water availability.

**Table 7 a.** Impact of water deficit on reproductive phase processes in Sitis.

| % of time process affected by water deficit |  |  |  |  |  |  |  |  |  |  |
| --- | --- | --- | --- | --- | --- | --- | --- | --- | --- | --- |
| SITIS |  |  |  |  |  |  |  |  |  |  |
| Genotypes | Panicle<br>primordium<br>formation | Flag leaf<br>formation | Lower<br>Internode<br>elongation | Branching I<br>formation | Branching II<br>formation | Internode 2<br>elongation | Spikelet<br>formation | Internode 1<br>elongation | Pollen<br>viability | Peduncle<br>elongation |
| CIRAD 409 | 0 | 0 | 0 | 0 | 0 | 0 | 73 | 89 | 100 | 100 |
| HD 1-4 | 0 | 0 | 0 | 23 | 33 | 33 | 73 | 100 | 100 | 100 |
| CIRAD 392 | 0 | 0 | 0 | 30 | 38 | 38 | 94 | 100 | 100 | 100 |
| IAC 25 | 0 | 0 | 0 | 30 | 38 | 38 | 94 | 100 | 100 | 100 |
| EM IAC 165 | 0 | 0 | 0 | 36 | 44 | 44 | 94 | 100 | 100 | 100 |
| GUARANI | 0 | 0 | 0 | 45 | 63 | 63 | 100 | 100 | 100 | 100 |

**Table 7 b.** Impact of water deficit on reproductive phase processes in Growth Chamber.

| % of time process affected by water deficit |  |  |  |  |  |  |  |  |  |  |
| --- | --- | --- | --- | --- | --- | --- | --- | --- | --- | --- |
| Growth chamber |  |  |  |  |  |  |  |  |  |  |
| Genotypes | Panicle<br>primordium<br>formation | Flag leaf<br>formation | Lower<br>internode<br>elongation | Branching I<br>formation | Branching II<br>formation | Internode 2<br>elongation | Spikelet<br>formation | Internode 1<br>elongation | Pollen<br>viability | Peduncle<br>elongation |
| IAC 25 | 0 | 38 | 38 | 100 | 100 | 100 | 100 | 100 | 100 | 100 |
| EM IAC 165 | 10 | 50 | 50 | 100 | 100 | 100 | 100 | 100 | 100 | 100 |
| CIRAD 392 | 23 | 62 | 62 | 100 | 100 | 100 | 100 | 100 | 100 | 100 |
| HD 1-4 | 44 | 69 | 69 | 100 | 100 | 100 | 100 | 100 | 100 | 100 |
| GUARANI | 57 | 79 | 79 | 100 | 100 | 100 | 100 | 100 | 100 | 100 |
| CIRAD 409 | 64 | 84 | 84 | 100 | 100 | 100 | 100 | 100 | 100 | 100 |

The percentages of the calculated relative progress of the reproductive phase are referring to the irrigated treatment and do not take into account any possible extension of the phase under water deficit.

**Table 8.** Analysis of variance of the flag leaf length in Sitis and Growth Chamber.

| SV | Sitis |  |  | Growth Chamber |  |  |
| --- | --- | --- | --- | --- | --- | --- |
|  | DF | Pr (>F) | Significance | DF | Pr (>F) | Significance |
| **Block** | 2 | 0.86 | NS | - | - | - |
| **Treatments** | 1 | 0.40 | NS | 1 | **0.00** | **\*\*\*** |
| **Genotypes** | 5 | 0.27 | NS | 5 | 0.30 | NS |
| **Trmnt. X Genotypes** | 5 | 0.22 | NS | 5 | 0.96 | NS |
| **CV%** |  | 11.79 |  |  | 19.36 |  |

Means were calculated on 6 genotypes x 3 replicates (18 values) for each water treatment in each experiment. NS: Not significant; \*\*\*: significant at $P < 0.001$

**Figure 5.**
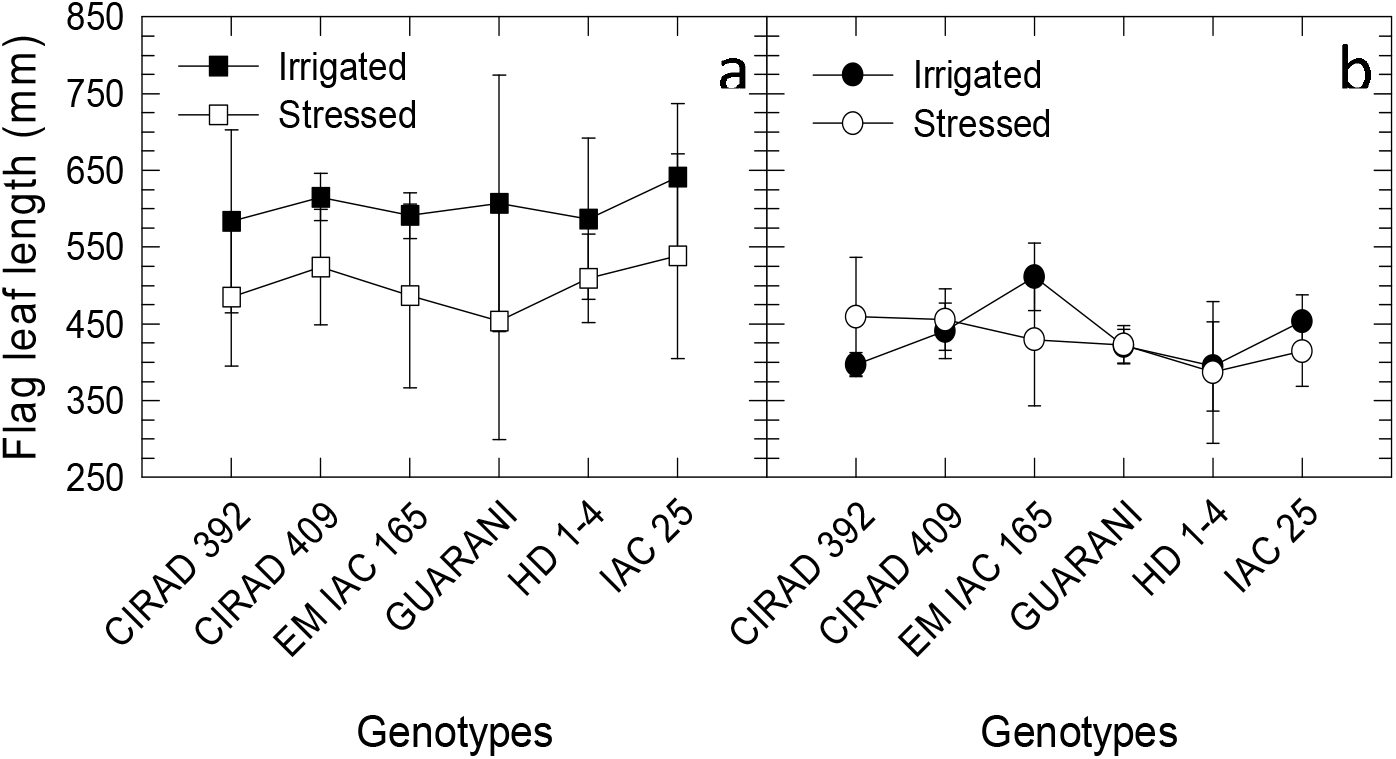
Flag leaf length in relation with the experiments and water treatments. Flag leaf length was measured at the end of experiment when plants were at heading. a) Growth chamber; b) Sitis

Three sets of relevant, discriminative and unbiased variables were selected in order to compare the genotypic responses to these water deficits. The first set concerned global indicators of development, growth and elongation measured during both treatments: TillerNb, Phyllochron, GrowthRate, ElongRate. The second set included key representative organs of the whole plant dimensioning and sink/source relationship: FLLength, FLWidth, IN2Length, IN2Width, and their associated sugar composition, FLSolSug, FLStarch, IN2SolSug and IN2Starch. The third set addressed the capacity of the plant to develop its reproductive structure driving the grain production: TotBranchNb and SpikeletNb. In addition, the transpiration rate (LeafTransp) was used as an indicator of the physiological status of the plant at the end of both treatments. Some traits were excluded because of the uncertainty on their process completion by the time when the last sampling was done. This was the case for IN1Length, IN1Width and PedLength. In fact, the whole trial was stopped at heading of the irrigated plants: at this time, plants under WD had not reached heading yet and their own progression to heading was variable, with high intra-variability for IN1 and Ped measurements. It was thus irrelevant to quantify the effect of water deficit on processes that were still on-going at the time of sampling, as any slight delay in development led to large differences in organ dimensions.

### Genotype response to water deficit at reproductive phase

The integrated response of the six genotypes to water deficit was analyzed based on the response indexes as defined in Material and Methods. The PCA of the selected morphological, physiological and biochemical traits was run using the variations of total branch number (ΔTotBranchNb) and spikelet number (ΔSpikeletNb) as supplementary variables. Indeed, considering the termination of the experiment at heading, these variables represent the overall performance of the plants as direct components of grain production and so were relevant indicators of plant adaptability to a WD established during the reproductive stage. The correlation matrix is given in Table 9. The supplementary variables ΔTotBranchNb and ΔSpikeletNb were highly correlated (+0.945***) to each other, as expected. The ΔTotBranch Nb was highly correlated to ΔFLLength (+0.730**), ΔFLWidth (+0.736**) and ΔIN2Width (+0.728**): the reduction in flag leaf and internode 2 dimensions in response to water deficit was closely associated with the reduction in the panicle branch number. The ΔSpikeletNb was also correlated to the same variables at 5% (*). The ΔLeafTransp was also positively correlated to ΔSpikeletNb (+0.578*) and logically more weakly to ΔTotBranchNb (+0.526°) as the formation of branches occurs prior to spikelet formation. In addition, ΔTillNb was the only trait negatively correlated to ΔTotBranchNb (−0.511°), and positively to ΔGrowthRate (+0.455°). The latter did not display any correlation with other traits, so was representative neither of organ growth (morphogenesis), or of organ generation (organogenesis). The ΔElongRate was positively correlated to ΔIN2Length (**) and ΔFLStarch (*), and negatively to ΔPhyllochron (**) and ΔFLSolSug (*). The ΔPhyllochron was positively correlated to ΔIN2SolSug.

**Table 9.** Correlation matrix of the studied variables.

| Variables | ΔGrowth rate | ΔElong rate | ΔTiller Nb | ΔFL length | ΔFL width | ΔIN2 length | ΔIN2 width | ΔLeafTransp | ΔFlagLeafSolSug | ΔFlagLeafStarch | ΔIN2SolSug | ΔIN2Starch | ΔPhyllochron | ΔTotalBranchNb |
| --- | --- | --- | --- | --- | --- | --- | --- | --- | --- | --- | --- | --- | --- | --- |
| ΔGrowthRate | 1 |  |  |  |  |  |  |  |  |  |  |  |  |  |
| ΔElongRate | 0.218 | 1 |  |  |  |  |  |  |  |  |  |  |  |  |
| ΔTillNb | 0.455° | -0.049 | 1 |  |  |  |  |  |  |  |  |  |  |  |
| ΔFLLength | -0.209 | -0.204 | -0.369 | 1 |  |  |  |  |  |  |  |  |  |  |
| ΔFLWidth | -0.196 | -0.054 | -0.521 | **0.580\*** | 1 |  |  |  |  |  |  |  |  |  |
| ΔIN2Length | 0.412 | **0.869\*\*\*** | -0.129 | -0.301 | -0.050 | 1 |  |  |  |  |  |  |  |  |
| ΔIN2Width | 0.149 | -0.001 | -0.498 | 0.435 | **0.712\*\*** | 0.266 | 1 |  |  |  |  |  |  |  |
| ΔLeafTransp | -0.188 | 0.291 | -0.419 | 0.131 | 0.044 | 0.438 | 0.396 | 1 |  |  |  |  |  |  |
| ΔFLSolSug | -0.188 | **-0.659\*** | -0.329 | 0.447 | 0.220 | **-0.594\*** | 0.239 | -0.194 | 1 |  |  |  |  |  |
| ΔFLStarch | 0.222 | **0.686\*** | -0.425 | -0.141 | 0.329 | **0.769\*\*** | 0.482 | 0.402 | -0.195 | 1 |  |  |  |  |
| ΔIN2SolSug | 0.287 | -0.518° | -0.009 | 0.151 | -0.052 | -0.209 | 0.458 | 0.097 | **0.593\*** | -0.055 | 1 |  |  |  |
| ΔIN2Starch | 0.203 | 0.453 | 0.218 | -0.254 | -0.322 | 0.434 | 0.063 | 0.217 | -0.221 | 0.370 | 0.305 | 1 |  |  |
| ΔPhyllochron | 0.045 | **-0.759\*\*** | -0.073 | -0.117 | -0.006 | -0.401 | 0.193 | -0.124 | 0.376 | -0.318 | **0.644\*** | -0.220 | 1 |  |
| ΔTotBranchNb | -0.178 | 0.047 | -0.511° | **0.730\*\*** | **0.736\*\*** | 0.032 | **0.728\*\*** | 0.526° | 0.207 | 0.375 | 0.193 | 0.025 | -0.136 | 1 |
| ΔSpikeletNb | -0.160 | 0.209 | -0.470 | **0.712\*\*** | **0.703\*** | 0.155 | **0.619\*** | **0.578\*** | 0.071 | 0.413 | -0.036 | -0.064 | -0.331 | **0.945\*\*\*** |

°:Significant at P< 0.1; \*: significant at P< 0.05; \*\*: significant at P< 0.01; and \*\*\*: significant at P< 0.001

The PCA representation on the two first components explained 55.4% of the total variability (Fig. 6). The first component (31.2%) was principally defined (i) positively by ΔElongRate and ΔIN2Length and (ii) negatively by ΔFLSolSug and ΔPhyllochoron. The second component (24.4%) was defined positively by ΔIN2Width and ΔFLWidth and negatively by ΔTillNb. The supplementary variables ΔTotBranchNb and ΔSpikeletNb were both positioned positively on the second component, and tightly associated with the maintenance of flag leaf width and internode 2 diameter (ΔFLWidth and ΔIN2Width). Tiller number variation was opposed to yield components variation on the main tiller and was the unique trait displaying negative correlations with them: thus, the more tillers were initiated under water deficit during the reproductive phase, the less branch and spikelet number was set in the main tiller panicle.

**Figure 6.**
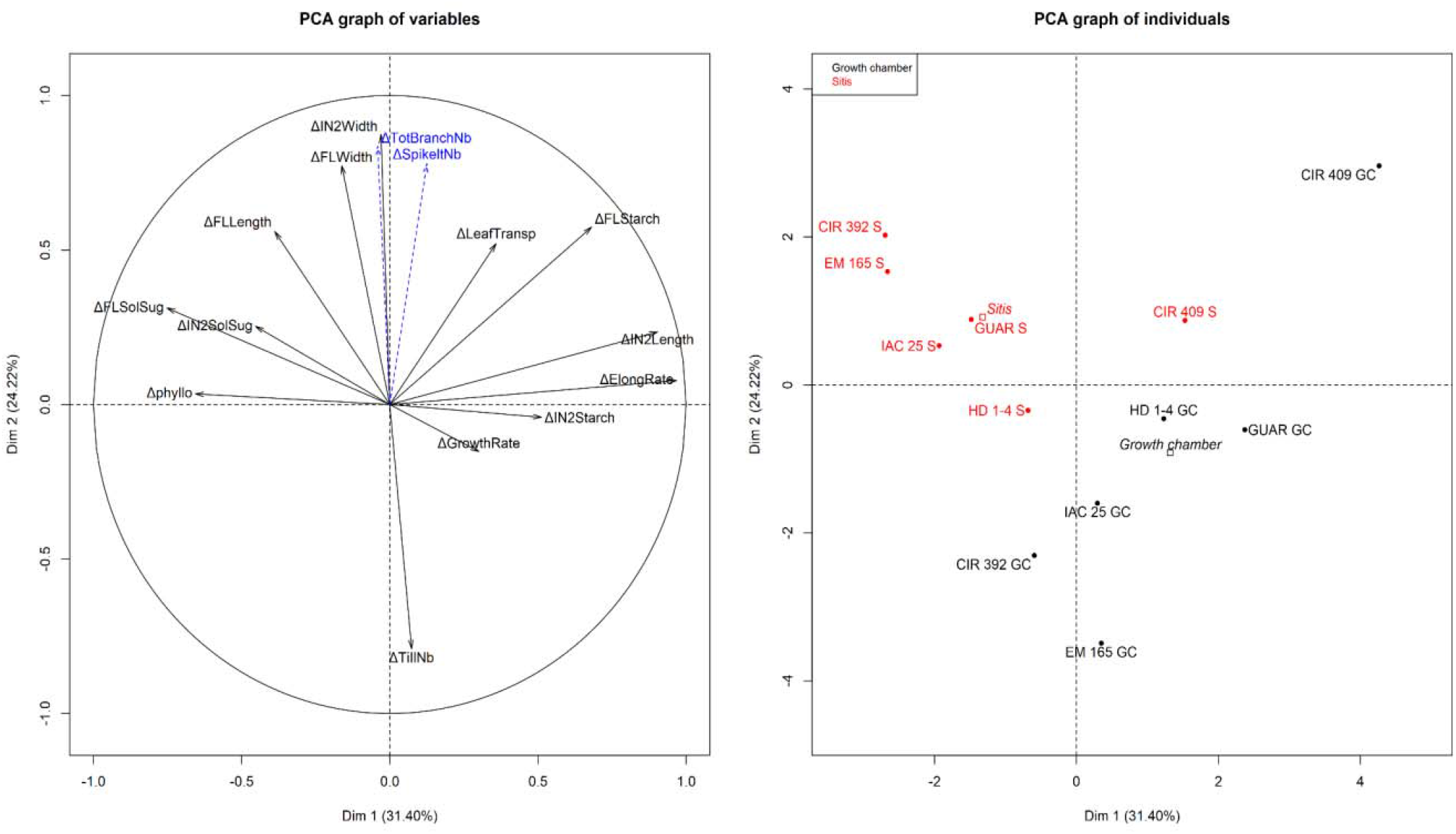
Principal Component Analysis representation on the two first components of the response index. PCA representation on the two first components, conducted on the response indexes in S and GC. ΔTotBranchNb and ΔSpikeltNb are supplementary variables, genotypes are supplementary observations. CIR 392 : Cirad 392; CIR 409 : Cirad 409; EM 165 : Early Mutant IAC 165; GUAR : Guarani; HD 1-4 : HD 1-4; IAC 25 : IAC 25. S : Sitis; GC : Growth Chamber

A large gradient of genotype responses was observed (Fig. 6 and Table 10) and opposed S and GC experiments. In both experiments, water deficit was associated with a reduction in the stem elongation (ΔElongRate, ΔIN2Length) and organ starch content (ΔFLStarch and ΔIN2Starch), while phyllochron and soluble sugar content increased (ΔFLSolSug and ΔIN2SolSug). This was illustrated by the genotype gradient along the PCA first component (Figure 6). In contrast, along the second component, the increase in soluble sugar was associated with the stability in yield components in S but not in GC.

**Table 10.** Response index in Sitis and in Growth Chamber experiments for all studied traits and genotypes.

|  | CIRAD 392 |  | CIRAD 409 |  | EM IAC165 |  | GUARANI |  | HD1-4 |  | IAC25 |  |
| --- | --- | --- | --- | --- | --- | --- | --- | --- | --- | --- | --- | --- |
| Traits | SITIS | Growth Chamber | SITIS | Growth Chamber | SITIS | Growth Chamber | SITIS | Growth Chamber | SITIS | Growth Chamber | SITIS | Growth Chamber |
| ΔTillNb | -0.12<sup>ns</sup> | 0.13<sup>ns</sup> | 0.04<sup>ns</sup> | -0.26<sup>ns</sup> | -0.10<sup>ns</sup> | 0.26<sup>ns</sup> | -0.13<sup>ns</sup> | 0.06<sup>ns</sup> | -0.07<sup>ns</sup> | -0.11<sup>ns</sup> | -0.27* | -0.06<sup>ns</sup> |
| ΔGrowthRate | -0.15 | -0.39** | -0.01<sup>ns</sup> | 0.03<sup>ns</sup> | -0.16<sup>ns</sup> | 0.41* | -0.19<sup>ns</sup> | -0.17<sup>ns</sup> | -0.21* | -0.38* | -0.38** | -0.21<sup>ns</sup> |
| ΔFLLength | 0.16<sup>ns</sup> | -0.17<sup>ns</sup> | 0.03<sup>ns</sup> | -0.15<sup>ns</sup> | -0.16<sup>ns</sup> | -0.25<sup>ns</sup> | 0.00 | -0.18<sup>ns</sup> | -0.02<sup>ns</sup> | -0.13<sup>ns</sup> | -0.09<sup>ns</sup> | -0.16<sup>ns</sup> |
| ΔFLWidth | 0.08<sup>ns</sup> | 0.00 | 0.04<sup>ns</sup> | 0.07<sup>ns</sup> | 0.03<sup>ns</sup> | -0.17** | -0.02<sup>ns</sup> | -0.12<sup>ns</sup> | -0.07<sup>ns</sup> | -0.11<sup>ns</sup> | -0.03<sup>ns</sup> | -0.05<sup>ns</sup> |
| ΔIN2Length | -0.39* | -0.34* | -0.04<sup>ns</sup> | 0.21<sup>ns</sup> | -0.32* | -0.12<sup>ns</sup> | -0.26* | -0.06<sup>ns</sup> | -0.27* | -0.13<sup>ns</sup> | -0.22* | -0.28** |
| ΔIN2Width | -0.03<sup>ns</sup> | -0.18** | 0.00 | 0.01<sup>ns</sup> | 0.03<sup>ns</sup> | -0.17** | -0.03<sup>ns</sup> | -0.09<sup>ns</sup> | -0.14* | -0.16*<sup>n</sup> | -0.09<sup>ns</sup> | 0.16** |
| ΔTotBranchNb | 0.09<sup>ns</sup> | -0.31* | -0.07<sup>ns</sup> | -0.08<sup>ns</sup> | -0.12<sup>ns</sup> | -0.46** | -0.13<sup>ns</sup> | -0.19<sup>ns</sup> | -0.23* | -0.17<sup>ns</sup> | -0.31** | -0.25<sup>ns</sup> |
| ΔSpikeletNb | 0.06<sup>ns</sup> | -0.28* | -0.01<sup>ns</sup> | -0.03<sup>ns</sup> | -0.17* | -0.45** | -0.19* | -0.24<sup>ns</sup> | -0.17* | -0.10<sup>ns</sup> | -0.33** | -0.26* |
| ΔElongRate | -0.44** | -0.34** | -0.20<sup>ns</sup> | -0.01<sup>ns</sup> | -0.55** | -0.33** | -0.38** | -0.17* | -0.31* | -0.26** | -0.41** | -0.28** |
| ΔFLSolSug | 0.90** | 0.26<sup>ns</sup> | -0.02<sup>ns</sup> | 0.18<sup>ns</sup> | 0.82** | 0.27* | 0.66** | 0.29* | 0.97** | 0.06<sup>ns</sup> | 0.68** | 0.15<sup>ns</sup> |
| ΔFLStarch | -0.18** | -0.43<sup>ns</sup> | -0.28** | 1.16<sup>ns</sup> | -0.13<sup>ns</sup> | -0.38<sup>ns</sup> | -0.34** | 0.14<sup>ns</sup> | -0.27** | -0.18<sup>ns</sup> | -0.39** | -0.37<sup>ns</sup> |
| ΔIN2SolSug | 1.48** | -0.62* | 0.08<sup>ns</sup> | -0.02<sup>ns</sup> | 1.91** | 1.21* | 1.10** | 1.40* | 0.38<sup>ns</sup> | -0.04<sup>ns</sup> | 0.85** | -0.35<sup>ns</sup> |
| ΔIN2Starch | -0.06<sup>ns</sup> | -0.34<sup>ns</sup> | -0.01<sup>ns</sup> | 0.17<sup>ns</sup> | -0.17<sup>ns</sup> | 0.09<sup>ns</sup> | -0.26** | 1.47<sup>ns</sup> | -0.17<sup>ns</sup> | -0.19<sup>ns</sup> | -0.20* | -0.06<sup>ns</sup> |
| ΔLeafTransp | -0.17<sup>ns</sup> | -0.36** | -0.05<sup>ns</sup> | -0.05<sup>ns</sup> | 0.06<sup>ns</sup> | -0.31* | -0.15<sup>ns</sup> | -0.05<sup>ns</sup> | -0.17* | 0.15<sup>ns</sup> | -0.20* | -0.25* |
| ΔPhyllochron | 0.25<sup>ns</sup> | 0.06<sup>ns</sup> | -0.03<sup>ns</sup> | -0.06<sup>ns</sup> | 0.53* | 0.32* | 0.18<sup>ns</sup> | 0.03<sup>ns</sup> | -0.14<sup>ns</sup> | 0.10<sup>ns</sup> | 0.45<sup>ns</sup> | 0.06<sup>ns</sup> |

Response index represents the relative variation of a trait between stressed and control treatments (see Mat 1 Meth); negative value means a reduction in stressed plants in relation with control ones. \*\*: significant by F test at 1% probability; \*: significant by F test at 5%; <sup>ns</sup>: not significant.

The singular performance of Cirad 409, minimizing its response to water deficit in both experiments, is noteworthy.

In S, no trait variation was observed with Cirad 409, except for flag leaf starch (ΔFLStarch). In contrast, high effects of water deficit were observed with the other genotypes. Flag leaf starch was reduced by 28%, while it was reduced by 39% with IAC 25 and 13% with EM IAC 165. As the most sensitive trait to WD, soluble sugars increased up to 97% in the flag leaf (HD 1-4) and to 148% in the internode 2 (Cirad 392). Main tiller elongation rate was also very sensitive to water deficit, with significant variations ranging between 31% (HD 1-4) and 55% (EM IAC 165). Variations of total branch number were smaller, mainly non-significant, except with IAC 25 and HD 1-4 (−31% and −23% respectively). Spikelet number was also poorly affected, except with IAC 25 (−33%), which highlighted the high response of IAC 25 to water deficit. Interestingly with Cirad 392, values of yield components were maintained under water deficit while significant variations in stem elongation and soluble sugar contents were reported.

In GC, no trait displayed significant variation with Cirad 409: all variables were maintained unchanged under moderate water deficit, as it was observed in S (with the exception of ΔFLStarch). The response to WD was quite different with other genotypes. Variation in flag leaf soluble sugars was much lower than in S, only Guarani and EM IAC 165 displayed a significant increase by +27 % and +29%, respectively. Oppositely, ΔIN2SolSug varied widely across genotypes: it decreased by 62% with Cirad 392, was stable with HD 1-4, and increased by 191% with EM IAC 165. The sensitivity of the elongation rate to water deficit was also remarkable with all genotypes except with Cirad 409: it decreased from 17% (Guarani) to 34% (Cirad 392). Finally, except for HD 1-4, yield components were more affected in GC than in S: EM IAC 165 and IAC 25 were the most sensitive genotypes, with a reduction in branch number by 46 and 25 % and spikelet number by 45 and 26%, respectively.

## Discussion

In this study, six rice genotypes were subjected to the same moderate water deficit during the reproductive phase by adjusting the pots to FTSW=0.4 every other day under two contrasted environments. In a Brazilian greenhouse experiment (S), water deficit started 15 days after panicle initiation and conditions were characterized with high temperature, high radiation and high VPD; in a French growth chamber experiment (GC), water deficit started 5 days after panicle initiation and conditions were characterized with optimum temperature, low radiation and low VPD.

### Contrasted climates generated different morphotypes

The vegetative and reproductive phases duration has been widely modified between both sites. The use of thermal time, instead of days, is widely accepted as a common way to estimate phase durations in varying environments. Underlying the assertion that development rate is driven by temperature (Jamieson et al. 1995), thermal time turns possible the comparison of plant phenology across environments despite their differences in temperature (Bouman et al. 2001). In the present study, not only temperature differed significantly across environments but also incoming radiation. Setting up an index combining both temperature and light variation could have been a relevant option to account for the variability in plant development. Indeed, to deal with environments differing in light as well, Islam and Morison (1992) introduced in rice a photothermal quotient (PQ) - as the ratio between cumulated radiation and cumulated thermal time (MJ.m^−2^.°Cdays^−1^) - during the 30 day period before harvest in order to predict yield. More recently, Baumont et al. (2019) indicated in wheat the joint role of temperature and carbon availability on plant development rate and concluded that the use of a photothermal quotient (PQ) is relevant to better model cycle durations in different light and temperature environments. In this study however, thermal time (TT) was relevant to account for development rate despite the contrasted light and temperature conditions: in the reproductive phase, period when the same number of organs emerged and developed in GC and S (the 4 last leaves and the panicle), thermal time was equivalent for both sites. In the vegetative phase, the larger amount of thermal time required to complete the phase in GC was fully explained by the extra number of developed phytomers: the plants produced an additional 1.5 leaf number for an extended vegetative period of 80 °C.days, which was equivalent to an overall mean phyllochron of 53 °C days, value in the range of those observed by Clerget et al., (2016).

The 63 % reduction in incoming radiation in GC compared to S, however, led to a highly significant decrease in tillering. Setting up a 70% reduction in incoming radiation in the field through nets, (Lafarge et al., 2010) reported a cessation in tiller emergence as soon as this was applied for 10 days at the vegetative stage, and a strong reduction in tiller emergence as soon as this was applied at the early reproductive phase. It also reduced the starch content in the flag leaf and stem internodes, and drastically decreased plant growth rate as reported by Lafarge et al (2010). In compensation, the lower temperature in GC resulted in a slower daily thermal time accumulation and in a consecutively major extension in days of the vegetative and reproductive phases. Overall, no significant differences were observed in plant biomass or internode length between sites, highlighting the existence of trade-offs between growth rate and duration, and between elongation rate and duration as well, in agreement with previous studies. In fact, Cookson et al. (2005) observed in leaf of *Arabidopsis thaliana* that initial expansion rate was negatively correlated with the duration of expansion. Also with *Arabidopsis thaliana* but using radiation treatments, Chenu et al. (2005) found an increase in duration along with a decrease in elongation rate under low radiation treatment, while Cookson and Granier (2006) observed that the trade-off was not complete. In contrast, a complete compensation between the extended duration and the decrease in elongation rate was reported by Aguirrezabal et al. (2006) when *Arabidopsis thaliana* was submitted to water deficit. In another side, the observed reduction in leaf size in S could be assigned to the high VPD effect on this site, as mentioned by Tardieu (2005) and Bouchabké et al (2006).

The development pattern of the genotypes under study has been modified across both sites when analyzed under favorable water conditions, but the plant biomass was globally unchanged, being set up in a longer time at a slower rate in GC. This difference in the time frame of the successive phenological events was crucial in analyzing the subsequent mild water deficit effect on organ growth and plant performance.

### A common physiological response to water deficit under both conditions led to different impacts on yield components

The reduction in elongation rate was observed on both experiments and was highly correlated to an increase in soluble sugars. The soluble sugar content in leaves is reported as increasing in response to water deficit in rice (Luquet et al, 2008; Rebolledo et al, 2012), because organ elongation is reduced before any effect on leaf photosynthesis is detected (Boyer 1970; Muller et al. 2011). Thus, the plant diminishing carbon demand leads to an accumulation of sugars in source leaf and sink organs, which in turn appears as a good indicator of stress perception. Similarly, the decrease in elongation rate was highly correlated to phyllochron lengthening, generating a delay in panicle emergence, a trait widely considered as indicative of plant sensitivity to water deficit (Garrity and O’Toole 1994; Boonjung and Fukai 1996b; Subashri et al. 2009; Sellamuthu et al. 2011; He and Serraj 2012; Sheoran et al. 2014; Swamy et al. 2017). Nevertheless, if the decrease in elongation rate looked related to yield potential loss (decrease in TotBranchNb and SpikeletNb) in GC, it was not the case in S (Fig.6): the more ElongRate decreased within the genotype set, the more TotBranchNb decreased in GC and increased in S. The maintenance of the stem elongation is therefore not systematically concomitant with the development of the reproductive organs. Overall, no correlation was observed between the two trait variations, meaning the adaptive value of the elongation rate response is dependent on plant growing conditions, and emphasizing how much conclusions often have to be restricted to the environment under study. Here, we hypothesize that plant C availability could explain the differences: C could become a limiting factor for growth and development under low radiation conditions in GC, while it was not in S.

### Some morphological traits were tightly linked to the maintenance of yield components across climate and water conditions

Oppositely, FLwidth and IN2Width reduction was tightly correlated to the reproductive sink size (branch and spikelet number) reduction across both environments. Also in rice Matsushima (1966) early highlighted a strong correlation between internode 1 thickness and spikelet number on the panicle. Likewise, Dingkuhn et al. (2015) and Adriani et al. (2016), working on a large panel of rice genotypes, observed a high positive correlation between flag leaf area and spikelet number on the panicle. With different nitrogen treatments, Kobayasi et al. (2002) demonstrated the correlation between the apical dome diameter at panicle initiation and the number of branches and spikelets. And when comparing wild (*Oryza Barthii*) and domesticated (*Oryza Glaberrima*) african rice, Ta et al (2017) reported in *O. Glaberrima* wider rachis meristem associated with a more branched panicle. In fact, correlations between apical meristem and leaf size were widely reported: according to Itoh et al. (2005), the rice apical meristem size increased with leaf rank, generating longer and wider leaves; in four Poa species Fiorani et al. (2000) observed that leaf size and elongation rate increased with meristem size; and Lacube et al. (2017) found in maize that meristem size drove leaf width. These results pinpoint that meristem size and activity conditioned organogenesis (branch and spikelet number) and morphogenesis (leaf dimension) of new generated organs. In our study, the maintenance of FLWidth and IN2Width was probably the phenotypic expression of the apical meristem size and activity maintenance, that in turn shall have preserved branch and spikelet formation under water deficit under both growing conditions.

A comprehensive representation of tolerance to water deficit during the reproductive phase relies therefore on the relationship between these trait variations (Fig. 7): the more FLWidth is reduced on both environments, the more TotBranchNb is reduced. A same tight correlation was obtained between variations in IN2Width and TotBranchNb (not shown). It also shows the consistency in the response across both sites of four genotypes, Cirad 409, Guarani, HD 1-4 and IAC 25, despite the contrasted conditions of incoming radiation and temperature, and suggests the validity of their responses over a large range of conditions. Oppositely, the response of the other two genotypes, EM IAC 165 and Cirad 392, was inconsistent over both experiments, because of their poor adaptation to specific conditions: Cirad 392, originating from the high Malagasy plateau is obviously not adapted to high temperature conditions in S, while EM IAC 165 cannot perform under low radiation in GC. Their tolerance classification changed from one place to the other, without affecting the correlation between traits. Overall, this study underlined the global tolerance of Cirad 409 and sensitivity of IAC 25.

**Figure 7.**
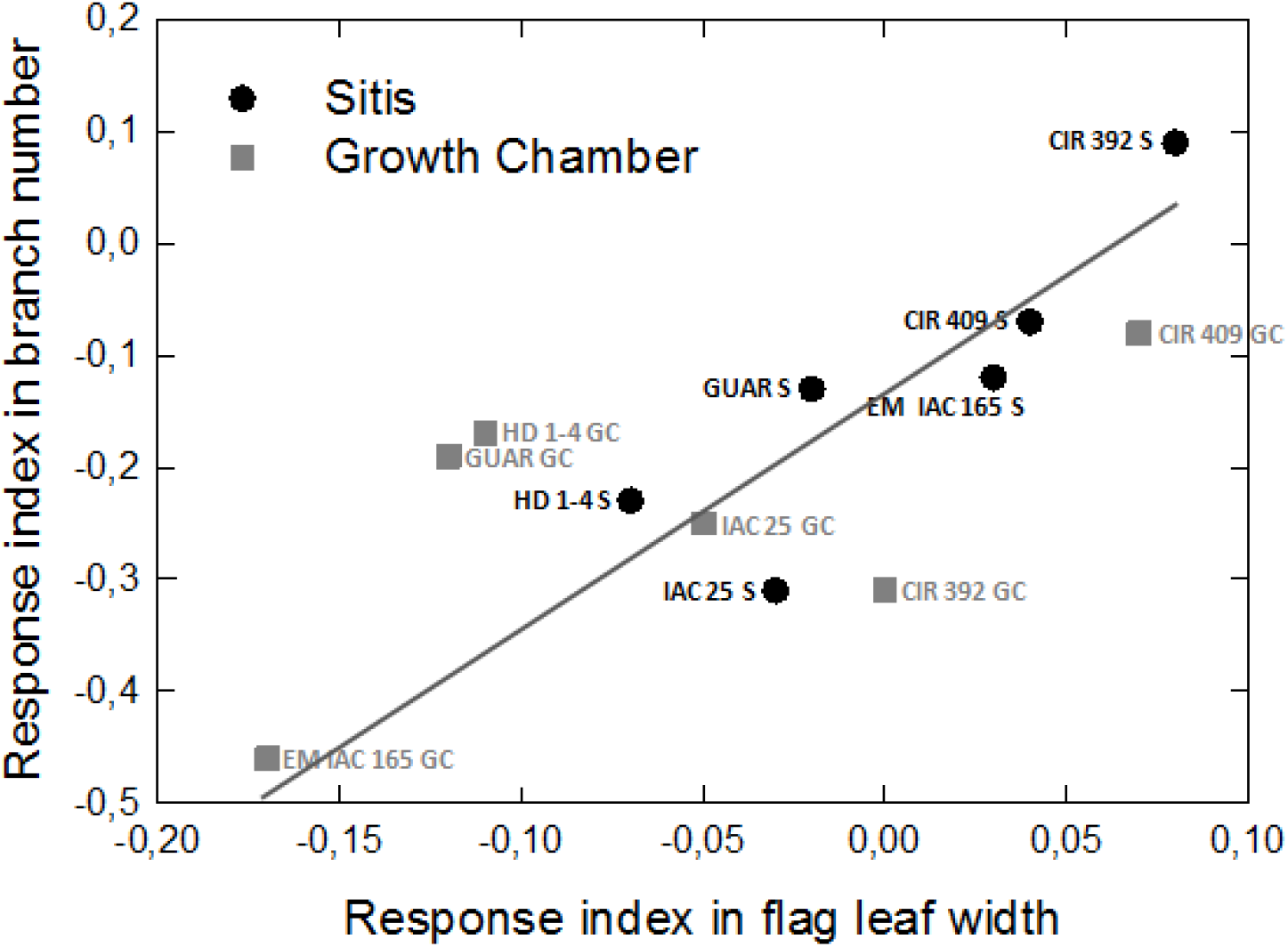
Relative variation of branch number and flag leaf width under water deficit. CIR 392 : Cirad 392; CIR 409 : Cirad 409; EM 165 : Early Mutant IAC 165; GUAR : Guarani; HD 1-4 : HD 1-4; IAC 25 : IAC 25. S : Sitis; GC : Growth Chamber

### Phenotyping for tolerance to mild water deficit at reproductive stage

The steady correlations obtained above do not allow comparing the genotype responses across both sites because of a lag time in water deficit application. Any classification of genotypes for mild water deficit tolerance would require applying the constraint at the same phenological age for all genotypes, more obviously when assessing the tolerance by the final yield and its components. It is therefore a main challenge for breeders. Indeed, when dealing with a wide phenotypic diversity, the application of a single water deficit period might affect genotypes at quite distinct stages, depending on their respective earliness. In order to minimize any discrepancy in phenological stage, several methods have been used in the literature. Garrity and O’Toole (1994) synchronized the flowering dates of 55 cultivars by clustering genotypes and by planting date respective of their cycle duration (based on previous data). In contrast, Lilley and Fukai (1994) triggered water deficit at a single fixed time after PI had occurred for each material. In the same line, Subashri et al. (2009) used the same date for establishing the water deficit accepting some slight differences in earliness within a set of near genetic materials. A precise and more time-consuming option implemented by Sellamuthu et al. (2011) is to regularly dissect extra plants and withhold irrigation when 50% of the studied genetic population is at PI, but with the high probability of not impacting the reproductive traits with the same intensity in all materials.

In the present study, the PI date was “a posteriori” determined in all genotypes by using the Haun index method. It made possible to quantify how much a basic process has been affected by the water deficit, and finally to assess the genotype responses on the only traits that were actually affected. This approach should be particularly useful in field conditions, within breeder’s trials or large-scale phenotyping trials, where the triggering of a water deficit cannot be driven at the plant or genotype level, as we did in controlled conditions with plants in pipes or in pots.

## Conclusions

The plants grown in the two contrasted environments under study displayed phenological and morphological differences at the time when water deficit was applied. Despite these distinct initial phenotypes, plants reacted the same way to water deficit: increase in phyllochron and in soluble sugar content in source and sink organs, that were associated with a decrease in stem elongation rate. Nevertheless, this conserved physiological response did not have the same impact on yield components across environments, highlighting the need to restrict the validity of results and interpretation to the conditions explored by the study.

One key methodological practice here during the reproductive phase was to identify the basic traits that were effectively affected by the transient water deficit -thanks to Haun index monitoring- and to conduct phenotyping only on these traits. In a first step, it was necessary to map panicle initiation respective of each genotype and to compare them on common affected traits. In a second step, the adaptive value of the trait response was assessed by measuring its impact on yield component maintenance and allowed to classify genotypes.

The study also explored the relationship between vegetative growth and reproductive organogenesis under water deficit, questioning whether they are synergistic or antagonistic. Are there prioritizing mechanisms for reproductive sink development in water deficit tolerant materials? At the main stem level, the study revealed a synergy through the maintenance of the apical meristem size. At the plant level, the study could not conclude but highlighted a weak negative correlation between the formation of new -and unfertile- tillers after panicle initiation and the maintenance of yield components. Further investigations are needed to clarify whether this putative prioritization of growth to the existing organs at the expense of new vegetative structures could be drought adaptive.

Overall, the flag leaf width maintenance had been pointed out as a key phenotypic trait to detect tolerance in case of a water deficit occurring at the early and medium stages of the reproductive phase. It shall express how much the plant is able to maintain its apical meristem size and activity and directly drives the maintenance in reproductive structure formation. In both environments tested in this study, flag leaf width reduction in response to water deficit was highly correlated with the reduction in branch and spikelet number in the panicle of the same tiller. We emphasize here that other traits could also be used for screening for drought tolerance according to the precise timing of water stress, like the variation in the diameter of an internode or the peduncle. The advantage to target such “secondary traits” resides in their capacity (i) to have a general value whatever the climate conditions; 2) to finely quantify the genotype-based effect of water deficit and so to distinguish the stress response across genotypes and (ii) to detect relevant QTLs, closer from the basic metabolic processes that drive rice plant response to water deficit.

## Declarations

### Ethics approval and consent to participate

Not applicable

### Consent for publication

Not applicable

### Availability of data and material

All data generated or analysed during this study are included in this published article.

### Authors’ contributions

APdC and TL coordinated the Dryce research project.IPdL, MdR, and APdC designed and carried out the Sitis experiment in Goiania. IPdL, SR, MdR and TL designed and carried out the growth chamber experiment in Montpellier. ACV and AS performed non-structural carbohydrate measurements and their statistical analysis. IPdL and MdR, with help of FB and TL, performed the data analysis and interpretation.MdR, IPdL and TL wrote the paper,which was edited and approved by all co-authors.

## Acknowledgements

The authors thank the Agropolis Foundation (France) and Embrapa (Brazil) for funding the bilateral DRYCE project and Capes (Brazil) for supporting Isabela’s PhD scholarship. The authors would like to thank all the Embrapa staff who contributed to the experiment, with particular thanks to Cleicomar Gonçalves de Almeida for help in conducting the experiment in the Sitis platform. Finally, we would like to thank Lauriane Rouan and Denis Cornet (Cirad) for help in the statistical analyses.

## List of abbreviations

TPE: Target populations of environment
HFE: Highly favorable environment
FE: Favorable environment
LFE: Least favorable environment
QTL: Quantitative trait loci
PI: Panicle initiation
SITIS: Integrated System for Induced Treatment of Drought
PVC: PolyVinyl Chloride
IRR: Fully irrigated
STR: Water stress
NPK: Nitrogen phosphorus ans potassium
FC: Field capacity
WP: Wilting point
FTSW: Fraction of transpirable soil water
T min: Minimal temperature
T max: Maximal temperature
Rad: Daily cumulated incident radiation
VPD: air vapor pressure deficit
Tb: Base temperature
Topt1: Temperature optimum 1
Topt2: Temperature optimum 2
HI: Haun Index
DAP: Days after planting
FLNb: Flag leaf number
RGR: Relative growth rate
GR: Growth rate
Exp.: Experiment
Trmnt: Treatment
IN: Internode
Max: Maximum
vs: versus
PCA: Principal component analysis
ΔGrowthRate: Relative variation of growth rate between IRR and ST treatments
ΔElongRate: Relative variation of total elongation rate between IRR and ST treatments
ΔTillNb: Relative variation of total tiller number between IRR and ST treatments
ΔFLLength: Relative variation of total flag leaf length between IRR and ST treatments
ΔFLWidth: Relative variation of total flag leaf width between IRR and ST treatments
ΔIN2Length: Relative variation of total internode 2 length between IRR and ST treatments
ΔIN2Width: Relative variation of total2 width between IRR and ST treatments
ΔLeafTransp: Relative variation of total leaf transpiration between IRR and ST treatments
ΔFLSolSug: Relative variation of total flag leaf soluble sugars between IRR and ST treatments
ΔFLStarch: Relative variation of total flag leaf starch between IRR and ST treatments
ΔIN2SolSug: Relative variation of total internode 2 soluble sugar between IRR and ST treatments
ΔIN2Starch: Relative variation of total internode 2 starch between IRR and ST treatments
ΔPhyllochron: Relative variation of total phyllochron between IRR and ST treatments
ΔTotalBranchNb: Relative variation of total branch number between IRR and ST treatments
ΔSpikeletNb: Relative variation of total spikelet number between IRR and ST treatments
TillNb: Tiller number
FLLength: Flag leaf length
FLWidth: Flag leaf width
IN2Length: Internode 2 length
IN2Width: internode 2 width
TotalBranchNb: Total branch number
SpikeletNb: Spikelet number
ElongRate: Elongation rate
FLSolSug: Flag leaf soluble sugars content
FLStarch: Flag leaf starch content
IN2SolSug: Internode 2 soluble sugars content
IN2Starch: Internode 2 starch content
LeafTransp: Leaf transpiration rate
PQ: Photothermal quotient

